# Flux analysis resolves distinct brain uptake and clearance of TfR- and CD98hc-targeting large molecules

**DOI:** 10.64898/2026.11.12.766279

**Authors:** Arash Moshkforoush, Darren Chan, Ahlam Qerqez, Jennifer E. Kung, Kylie S. Chew, Robert C. Wells, Johann Chow, Richard Theolis, Nicholas P.D. Liau, Joseph Duque, Padma Akkapeddi, Abira Bandyopadhyay, Isabel Becerra, Butch Benitez, Ellen Y. T. Chien, Roni Chau, Allisa Clemens, David Huynh, Do Jin Kim, Nathalie Khoury, Kendra J. Lechtenberg, Eric Liang, Hiwot Misker, Hoang Nguyen, Michelle E. Pizzo, Kevin Rebadulla, Yaneth Robles-Colmenares, Mahdiar Sadeghi, Tiffany Tran, Ranya K.A. Virk, Kaitlin Xa, Annie Arguello, Mark S. Dennis, Joseph W. Lewcock, Cathal S. Mahon, Lionel Rougé, John Michael Sauer, Ryan J. Watts, Don Zhong, Y. Joy Yu Zuchero, Kapil Gadkar, Mihalis S. Kariolis, Shyeilla Dhuria, Robert G. Thorne, Christopher M. Koth, Kirk R. Henne

## Abstract

Advances in large-molecule delivery across the blood-brain barrier (BBB) by targeting receptors such as transferrin receptor (TfR) or CD98 heavy chain (CD98hc) are shifting the focus from restricted parenchymal access toward brain exposure profiles tailored to therapeutic needs. However, brain concentrations offer only partial insight into delivery mechanisms, as they reflect the net balance of two dynamic processes: flux of molecules entering and leaving the brain, here termed *brain uptake* and *brain clearance*, respectively. Here, we establish a generalizable experimental and mathematical modeling framework to resolve these opposing fluxes and demonstrate its application using pharmacokinetic (PK) data from Fc-engineered antibody transport vehicles (ATVs) targeting TfR or CD98hc in humanized knock-in mouse models. First, to directly estimate brain clearance, we engineered anti-ATV antibodies that selectively bind the TV patch to block receptor-mediated brain uptake *in vivo.* Integrating these measurements with a mechanistic PK model, we then reconstructed brain fluxes of ATV molecules over time across a broad range of receptor affinities. Results demonstrated that TfR-targeting promotes rapid, affinity-dependent brain uptake accompanied by comparatively rapid brain clearance, whereas CD98hc-targeting results in slower brain uptake and prolonged retention. Extending this framework to dual TfR/CD98hc-targeting ATVs suggested that higher observed brain exposure relative to single-receptor formats does not necessarily translate to greater brain delivery: increased exposure was predicted to arise primarily from reduced brain clearance, associated with CD98hc-mediated retention, rather than enhanced uptake through dual-receptor engagement. Together, these results establish a quantitative basis for resolving brain fluxes and engineering brain-targeted biologics for desired therapeutic outcomes.

## Introduction

The blood-brain barrier (BBB) restricts parenchymal access of systemically administered antibodies, presenting a major challenge for the treatment of central nervous system (CNS) disorders (1–4). Despite decades of clinical development and numerous trials, only a few biologics lacking active BBB-targeting mechanisms have demonstrated meaningful clinical efficacy (5–7). However, advances in receptor-mediated transcytosis (RMT), achieved by targeting highly expressed receptors at the BBB, have expanded opportunities for brain delivery of large-molecule therapeutics. These include the well-established transferrin receptor (TfR) and the more recently characterized CD98 heavy chain (CD98hc) (8–16). In particular, TfR-targeting has emerged as a strategy to enhance CNS delivery, with next-generation anti-amyloid antibodies, RNA interference therapeutics, and enzyme replacement therapies (ERTs) having advanced into clinical development (17–23). We have also previously reported the development of the transport vehicle (TV) technology, an antibody-based construct leveraging Fc-domain mutations that enable RMT by targeting TfR (4), CD98hc (24), or both receptors within a single molecule (25). The TV platform has demonstrated enhanced brain exposure compared to the control human IgG (huIgG) in preclinical animal models (4, 24, 26–32), with several TfR-targeting TV therapeutics having progressed to clinical evaluation or approval (33–35).

Disease biology and therapeutic mechanism of action (MOA) determine the optimal brain biodistribution and pharmacokinetics required to drive efficacy. For example, therapeutics requiring intracellular delivery, such as enzymes, oligonucleotides, and targeted protein degraders, must efficiently cross the BBB and reach endosomal or nuclear compartments (26, 27, 29, 32, 36, 37). In contrast, therapies targeting membrane-bound proteins or extracellular species are more likely to benefit from prolonged parenchymal retention and sustained target engagement (38). Within a therapeutic class, tissue biodistribution can also influence clinical outcomes. For instance, anti-amyloid antibodies with high vascular exposure have been linked to a high incidence of amyloid-related imaging abnormalities (ARIA), while receptor-targeted formats such as ATV.TfR can mitigate ARIA incidence by shifting antibody distribution toward the parenchyma (31).

In preclinical models, brain PK of biologics is typically assessed by measuring total drug concentrations in brain homogenates. However, these concentrations provide only a limited insight into the mechanisms underlying brain delivery, as they represent the net balance between two dynamic processes: the flux of molecules entering and leaving the brain, which, for simplicity, we refer to as *brain uptake* and *brain clearance* (Fig. 1A). Brain uptake includes passive or non-specific entry, primarily via the blood-CSF barrier, as well as active receptor-mediated transcytosis (RMT) (14, 15, 39–42). Conversely, brain clearance encompasses meningeal and perivascular pathways (43–45), cellular uptake followed by lysosomal degradation (catabolism) (4, 24, 46), or re-entry into systemic circulation via efflux transporters and bulk flow mechanisms (47).

**Fig. 1.**
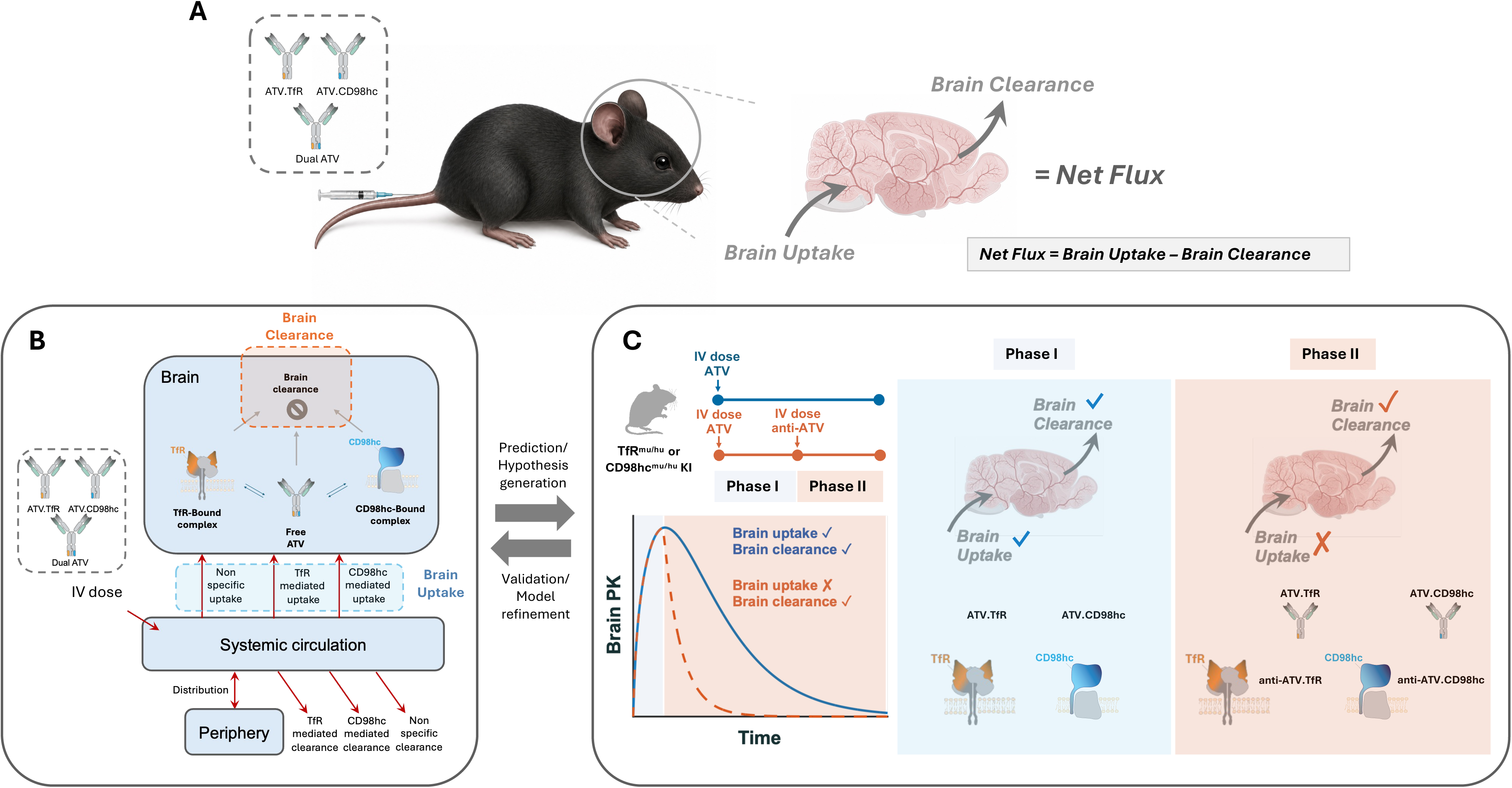
Conceptual framework for flux-based analysis of brain uptake and clearance of ATV molecules. **(A)** Schematic illustrating the dynamic balance between brain uptake and brain clearance that governs the brain PK of ATVs targeting TfR (ATV.TfR), CD98hc (ATV.CD98hc), or both receptors within a single dual ATV molecule following an intravenous administration. Brain uptake comprises both receptor-mediated (TfR- and CD98hc-mediated) and non-specific transport into the brain. Brain clearance represents the combined loss of ATV from the brain through receptor-mediated and non-specific elimination pathways. The net change in brain concentration is determined by the balance between brain uptake and brain clearance (net brain flux = brain uptake − brain clearance). **(B)** Overview of the mathematical modeling framework used to predict brain uptake and brain clearance fluxes of ATV molecules from the collective plasma and brain PK data. **(C)** Schematic of the pulse–chase strategy used to experimentally isolate the brain clearance. In the control arm (blue), humanized mice receive a single intravenous dose of ATV, where the brain PK reflect the combined effects of ongoing brain uptake and brain clearance. In the pulse–chase arm (orange), mice receive an intravenous dose of ATV followed by systemic administration of an anti-ATV antibody at the predicted brain T_max_. During Phase I, brain PK reflects concurrent brain uptake and clearance. Following anti-ATV administration (Phase II), receptor-mediated brain uptake is blocked while brain clearance is maintained, such that the subsequent decline in brain concentrations primarily reflects the brain clearance. Representative mouse and brain illustrations were AI generated. Brain anatomy and vasculature are schematic and shown for illustrative purposes only.

The same brain concentration at a given time point can result from very different combinations of uptake and clearance, obscuring the underlying transport kinetics. Additionally, conventional PK metrics such as maximal brain concentration (C_max_) and area under the brain concentration–time curve (AUC) cannot distinguish greater cumulative brain uptake from prolonged retention, limiting their ability to inform the rational design of brain-targeted therapeutics. Recent studies have further highlighted this limitation. For example, Veire et al. (46) used comparative radiotracing to investigate cellular uptake and catabolism of TfR- and CD98hc-targeting antibodies, revealing their distinct intracellular processing and brain retention properties.

Mathematical modeling provides a quantitative approach to deconvolute the underlying brain uptake and clearance fluxes from observed brain PK profiles (Fig. 1B). However, experimental measurements are essential to validate and refine model-based predictions. Because brain concentrations reflect the net balance between brain uptake and clearance, selectively suppressing one process provides a means to directly estimate the other. For example, blocking additional brain uptake once molecules have entered the brain enables estimation of their clearance without the confounding contribution of ongoing uptake. Conversely, minimizing brain clearance, e.g., by targeting brain-retentive epitopes such as myelin oligodendrocyte glycoprotein (MOG), could facilitate direct estimation of brain uptake (48). Together, this integrated modeling and experimental framework provides a generalizable strategy to better understand and quantify brain fluxes of brain-penetrant antibodies and assess how they are influenced by factors such as receptor binding affinity, dose level, and transport capacity.

Here, we demonstrate an application of this framework using ATV.TfR and ATV.CD98hc antibodies in mice expressing humanized TfR and CD98hc, i.e., TfR^mu/hu^ and CD98hc^mu/hu^ knock- in (KI), respectively (4, 24). We implemented the uptake-blocking strategy by engineering anti-ATV antibodies that selectively bind the TV patch with high affinity, preventing subsequent engagement of ATVs with TfR or CD98hc at the BBB, thereby blocking the additional brain uptake and enabling selective estimation of brain clearance (Fig. 1C). Integrating these measurements with the net brain flux, obtained from model-simulated brain PK profiles without anti-ATV intervention, allowed reconstruction of brain uptake over time for ATV.TfR and ATV.CD98hc across a broad range of receptor binding affinities. We then extended the flux analysis to dual ATVs engaging both TfR and CD98hc within a single molecule. Specifically, we investigated whether the transport properties of individual receptor-targeting ATVs could explain the enhanced brain exposure of dual ATVs relative to their corresponding single-receptor ATV formats (25). We also investigated whether this enhancement resulted from greater cumulative brain uptake, reduced brain clearance, or a combination of both. To isolate receptor-mediated transport from potential effects of therapeutic target binding, all ATV constructs were paired with Fabs targeting dinitrophenol (DNP), a small-molecule hapten absent in vivo (49).

## Results

### Distinct pharmacokinetic properties of ATV.TfR and ATV.CD98hc antibodies

The impact of receptor binding affinity on plasma and brain PK profiles was examined for ATV.TfR and ATV.CD98hc variants in TfR^mu/hu^ and CD98hc^mu/hu^ KI mouse models, respectively (Fig. 2 A-D). Following a single intravenous (IV) administration, both formats exhibited higher systemic clearance (CL) compared to an isotype-matched human IgG control (anti-DNP), with CL generally increasing in a receptor binding affinity–dependent manner (Fig. 2A and B; Table 1). This is consistent with target-mediated drug disposition (TMDD) (50), where TfR or CD98hc engagement in peripheral tissues increases systemic disposition of ATV molecules. At the administered dose of 50 mg/kg, ATV.TfR variants displayed ∼2-4x higher systemic CL than their ATV.CD98hc counterparts at comparable receptor binding affinities. Additionally, systemic CL was more sensitive to changes in receptor binding affinity for ATV.TfR than for ATV.CD98hc, with ∼5x difference across the tested TfR affinity range (K_D_ = 110-8000 nM), compared with ∼2.5x across the CD98hc affinity range (K_D_ = 170-6300 nM; Table 1).

**Fig. 2.**
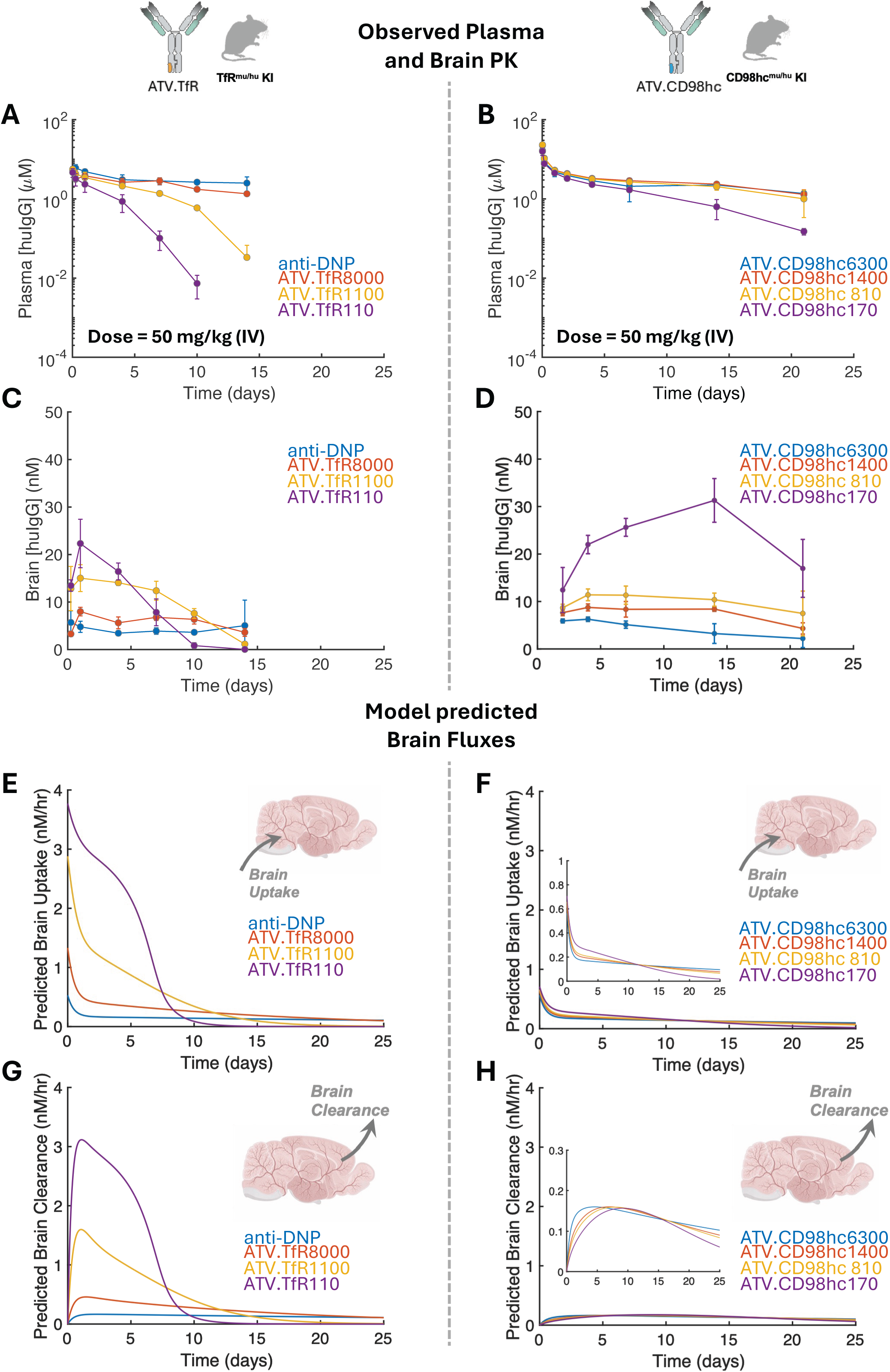
Observed systemic and brain PK and model-predicted brain fluxes of ATV.TfR and ATV.CD98hc in humanized mouse models. **(A–B)** Representative plasma PK profiles of ATV.TfR (A) and ATV.CD98hc (B) variants in TfR^mu/hu^ KI and CD98hc^mu/hu^ KI mice following a single 50 mg/kg IV dose, respectively. **(C–D)** Corresponding brain PK profiles of ATV.TfR (C) and ATV.CD98hc (D) variants. Observed data are shown as mean ± standard deviation (n = 4-5 animals/group/time point). **(E–H)** Model-predicted brain uptake **(E,F)** and clearance **(G,H)** fluxes for the corresponding ATV.TfR and ATV.CD98hc variants based on simulations from the calibrated PK model shown in Fig. 1B.

**Table 1.**
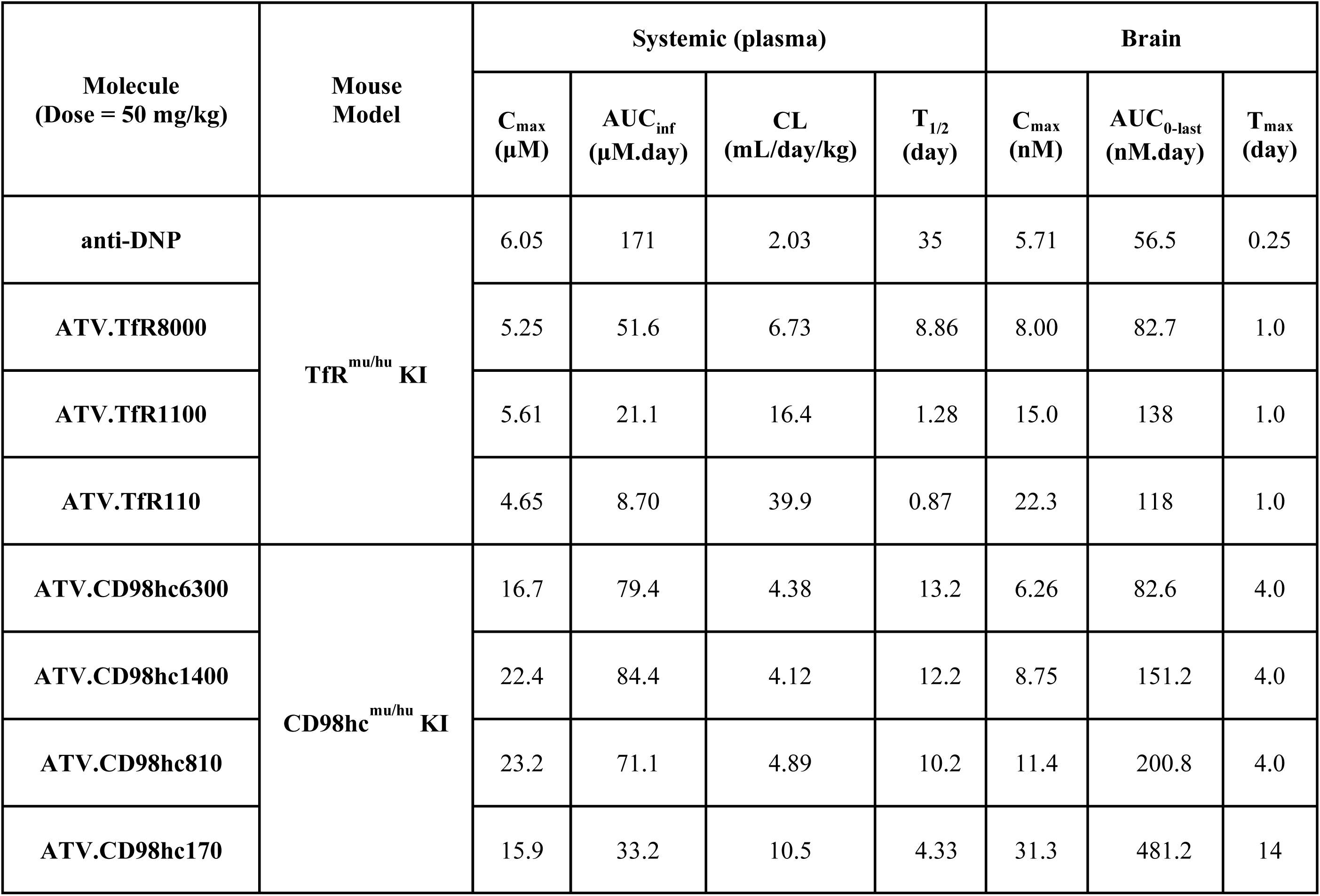
Summary of plasma and brain PK parameters for ATV variants corresponding to Fig. 2 A–D.

| **Molecule<br>(Dose = 50 mg/kg)** | **Mouse<br>Model** | **Systemic (plasma)** |  |  |  | **Brain** |  |  |
|---|---|---|---|---|---|---|---|---|
|  |  | **C<sub>max</sub><br>(µM)** | **AUC<sub>inf</sub><br>(µM.day)** | **CL<br>(mL/day/kg)** | **T<sub>1/2</sub><br>(day)** | **C<sub>max</sub><br>(nM)** | **AUC<sub>0-last</sub><br>(nM.day)** | **T<sub>max</sub><br>(day)** |
| **anti-DNP** | **TfR<sup>mu/hu</sup> KI** | 6.05 | 171 | 2.03 | 35 | 5.71 | 56.5 | 0.25 |
| **ATV.TfR8000** |  | 5.25 | 51.6 | 6.73 | 8.86 | 8.00 | 82.7 | 1.0 |
| **ATV.TfR1100** |  | 5.61 | 21.1 | 16.4 | 1.28 | 15.0 | 138 | 1.0 |
| **ATV.TfR110** |  | 4.65 | 8.70 | 39.9 | 0.87 | 22.3 | 118 | 1.0 |
| **ATV.CD98hc6300** | **CD98hc<sup>mu/hu</sup> KI** | 16.7 | 79.4 | 4.38 | 13.2 | 6.26 | 82.6 | 4.0 |
| **ATV.CD98hc1400** |  | 22.4 | 84.4 | 4.12 | 12.2 | 8.75 | 151.2 | 4.0 |
| **ATV.CD98hc810** |  | 23.2 | 71.1 | 4.89 | 10.2 | 11.4 | 200.8 | 4.0 |
| **ATV.CD98hc170** |  | 15.9 | 33.2 | 10.5 | 4.33 | 31.3 | 481.2 | 14 |

Both ATV.TfR and ATV.CD98hc variants exhibited higher brain concentrations relative to the isotype control (Fig. 2 C and D; Table 1), with C_max_ increasing with stronger receptor-binding affinity. However, their kinetic profiles differed markedly. ATV.TfR variants reached brain C_max_ rapidly, with a time to C_max_ (T_max_) of ∼1-day post-dose, followed by a decline in brain concentrations to below those of the isotype control within 10–15 days (Fig. 2C). Notably, the post-C_max_ decline was steeper with stronger TfR-binding affinity. In contrast, ATV.CD98hc exhibited a delayed T_max_ (≥ 4 days) and sustained brain exposures throughout the 21-day study across all affinity variants (Fig. 2D, Table 1).

### Model-predicted receptor-specific differences in brain fluxes of ATVs

To elucidate potential mechanisms underlying the observed differences in the brain PK of ATV.TfR and ATV.CD98hc, the PK model shown in Fig. 1B was calibrated to the pooled ATV data in humanized mouse models across a wide range of receptor binding affinities and dose levels (Materials and Methods; *SI Appendix,* Tables S1-4; *SI Appendix,* Figs. S1-3), and was used to simulate brain fluxes for the affinity variants across both ATV formats (Fig. 2 E-H). Simulations predicted a greater magnitude of brain uptake for ATVs relative to the isotype-matched control, consistent with the additional route of brain delivery via RMT. However, predicted brain uptake was substantially higher for ATV.TfR variants than ATV.CD98hc, with ∼5x difference in the peak uptake flux for the strongest tested affinities (Fig. 2 E and F). This difference, however, is counterbalanced by markedly higher predicted brain clearance for ATV.TfR (Fig. 2 G and H), resulting in a comparable brain C_max_ between the two formats despite pronounced differences in the kinetics and overall brain PK profiles. The model also provides a plausible mechanism for the earlier T_max_ observed with ATV.TfR (i.e., when net flux is zero: the point at which brain uptake and clearance intersect), as these molecules display faster transport kinetics for both brain uptake and clearance compared to the ATV.CD98hc counterparts (*SI Appendix,* Fig. S4).

### Engineering anti-ATV antibodies to selectively block receptor-mediated brain uptake

To directly estimate brain clearance without the confounding contribution of continued uptake, we developed an experimental approach to selectively block the receptor-mediated brain uptake of ATV.TfR and ATV.CD98hc molecules *in vivo*. Specifically, we developed mouse-derived anti-ATV antibodies that bind with strong affinity to ATV.TfR and ATV.CD98hc to prevent their binding to TfR or CD98hc receptors (Fig. 3). The anti-ATV antibodies demonstrated high-affinity binding to their respective ATV-modified Fc regions with no cross-reactivity to the wild-type human IgG1 Fc (*SI Appendix,* Table S6, *SI Appendix,* Fig. S5). High resolution structures of complexes of both anti-ATV Fabs with their respective monovalent Fc-only ATVs confirmed that both anti-ATV antibodies bind epitopes that overlap with their respective TfR- or CD98hc-binding interfaces on the Fc (Fig. 3A and B, *SI Appendix,* Fig. S6 A-D). Functional *in vitro* assays, including surface plasmon resonance (SPR), as well as cell-based competition assays in HEK cells, demonstrated that the anti-ATV antibodies effectively inhibit cellular binding of ATV.TfR and ATV.CD98hc variants (Fig. 3 C and D and *SI Appendix,* Fig. S6 E and F, respectively). The anti-ATV antibodies were subsequently reformatted to rabbit IgG (rbIgG) isotype and administered intravenously in WT C57/BL6 mice (10 mg/kg). The plasma PK profiles of anti-ATV.TfR and anti-ATV.CD98hc were highly similar, with an expected systemic CL of ∼7-8 mL/day/kg (*SI Appendix,* Fig. S6 G).

**Fig. 3.**
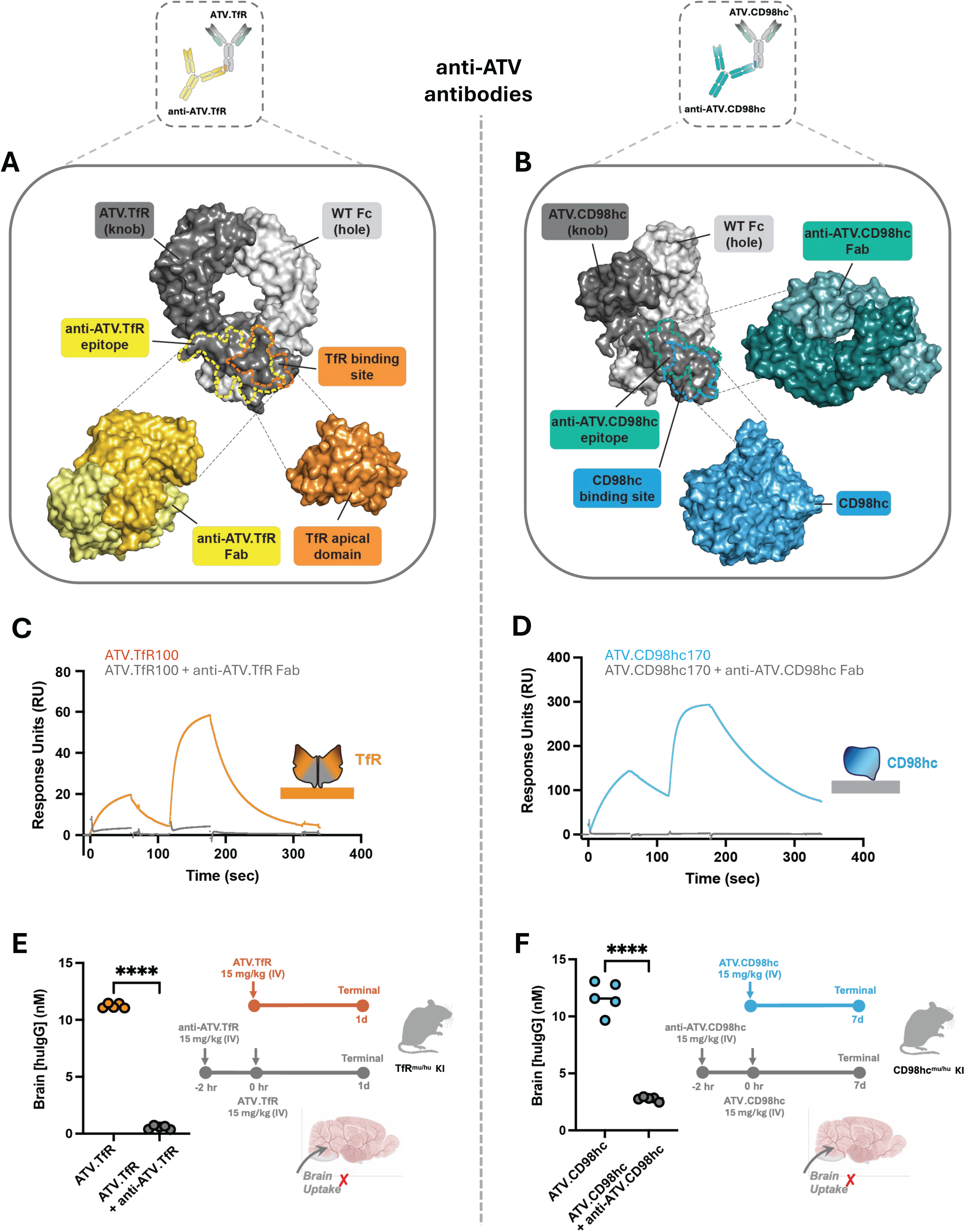
Anti-ATV antibodies compete with TfR and CD98hc for TV binding. **(A)** Surface representation of ATV.TfR in gray with the epitope of anti-ATV.TfR outlined in yellow and the binding interface of apical TfR outlined in orange. Surface representations of anti-ATV.TfR Fab (yellow) and TfR apical domain (orange) are shown below ATV.TfR in an open book representation, showing the relative binding orientations of both proteins on ATV.TfR. The binding interface and structure of the ATV.TfR-TfR apical domain interaction is adapted from PDB: 6W3H. **(B)** Surface representation of ATV.CD98hc in gray with the epitope of anti-ATV.CD98hc outlined in teal and the binding interface of CD98hc outlined in blue. Surface representations of anti-ATV.CD98hc Fab (teal) and CD98hc (blue) are shown below ATV.CD98hc in an open book representation, showing the relative binding orientations of both proteins on ATV.CD98hc. The binding interface and structure of the ATV.CD98hc-CD98hc interaction is adapted from PDB: 8G0M. **(C)** Surface plasmon resonance (SPR) demonstrating binding to two concentrations (333nM and 1µM, respectively) of ATV.TfR to immobilized biotinylated huTfR^ECD^ (orange) and blockade of binding in the presence of 3µM anti-ATV.TfR Fab (yellow). **(D)** SPR binding to two concentrations (333nM and 1µM, respectively) of ATV.CD98hc to immobilized biotinylated huCD98hc (blue) and blockade of binding in the presence of 3uM anti-ATV.CD98hc Fab (cyan). **(E)** TfR^mu/hu^ KI mice were dosed with 15mg/kg anti-ATV.TfR followed by injection of 15mg/kg ATV.TfR two hours later. Mouse brains were harvested 1 day later. Brain human IgG concentrations were measured using MSD. n=5 animals per group. **(F)** CD98hc^mu/hu^ KI mice were dosed with 15mg/kg rabbit anti-ATV.CD98hc followed by injection of 15mg/kg ATV.CD98hc two hours later. Mouse brains were harvested 7 days later. Brain human IgG concentrations were measured using MSD. n=5 animals per group. Statistical analysis was performed using an unpaired t-test (****p < 0.0001, ns = non-significant).

To validate the ability of anti-ATV antibodies to block brain uptake of ATV molecules *in vivo*, TfR^mu/hu^ KI and CD98hc^mu/hu^ KI mice were then pre-dosed with 15 mg/kg of rabbit anti-ATV.TfR or anti-ATV.CD98hc antibodies, respectively, followed two hours later by an IV administration of an equimolar dose of ATV.TfR or ATV.CD98hc (Fig. 3 E and F). Brain concentrations were assessed 1 day after ATV.TfR dosing and 7 days post ATV.CD98hc administration, consistent with their respective predicted brain T_max_ (*SI Appendix,* Fig. S1). In both models, pre-dosing with anti-ATV antibodies markedly reduced brain concentrations of the corresponding ATV molecule compared to non-pre-dosed controls, with reductions of >90% for ATV.TfR (Fig. 3E) and ∼80% for ATV.CD98hc (Fig. 3F). Collectively, these results demonstrate the feasibility of the anti-ATV strategy in inhibiting receptor-mediated brain uptake of systemically administered ATVs.

### Antibody-blocking “Pulse-chase” strategy enables quantification of brain clearance of ATV molecules

We next designed a “pulse-chase” experiment to directly and selectively measure the brain clearance of ATV.TfR and ATV.CD98hc molecules *in vivo* (Fig. 4). TfR^mu/hu^ KI and CD98hc^mu/hu^ KI mice were dosed with ATV.TfR or ATV.CD98hc, respectively, followed by systemic administration of an equimolar corresponding anti-ATV antibody at their respective predicted brain T_max_ (1 day for ATV.TfR and 7 days for ATV.CD98hc). By blocking further receptor-mediated brain uptake, this approach enabled selective characterization of the brain clearance of ATV molecules already present in the brain tissue. Three affinity variants from each ATV platform were tested (TfR K_D_ = 100, 620, and 2500 nM for ATV.TfR, and CD98hc K_D_ = 170, 550, and 6300 nM for ATV.CD98h) at a dose of 30 mg/kg to evaluate how receptor binding affinity influences kinetics of brain clearance.

**Fig. 4.**
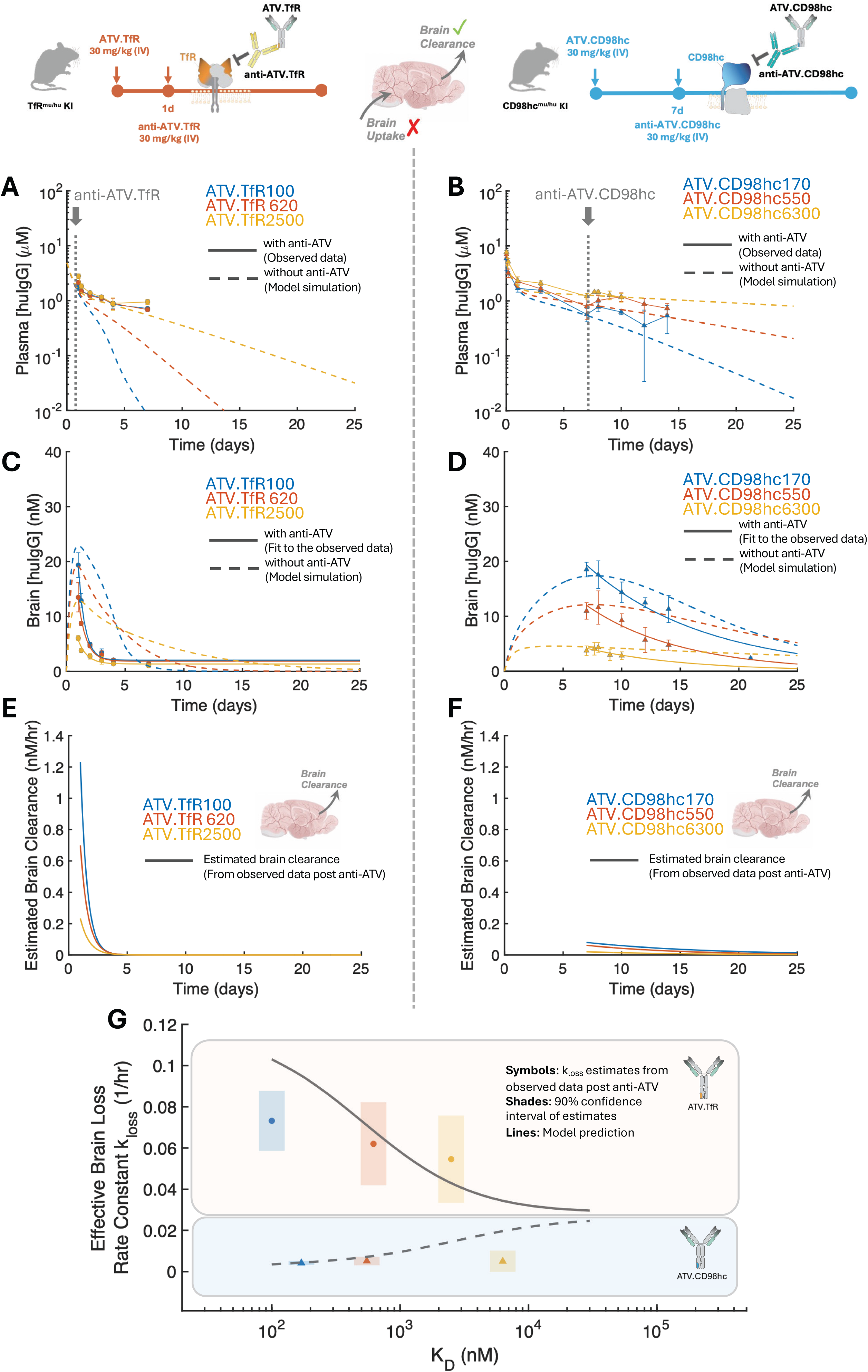
Experimental estimation of brain clearance of ATV.TfR and ATV.CD98hc using the pulse–chase strategy. **(A,B)** Plasma PK profiles of ATV.TfR (A) and ATV.CD98hc (B) affinity variants following a single 30 mg/kg intravenous dose followed by anti-ATV administration at the indicated time points. Symbols and solid traces represent the observed data, while dashed traces represent model-predicted PK profiles in the absence of anti-ATV administration. **(C,D)** Corresponding brain PK profiles of ATV.TfR (C) and ATV.CD98hc (D) variants. Solid traces represent mono-exponential fits to the observed post-anti-ATV brain PK profiles (symbols), and dashed traces represent model-predicted brain PK profiles without anti-ATV administration. **(E,F)** Brain clearance fluxes for ATV.TfR (E) and ATV.CD98hc (F) variants estimated from a time derivative of the post-anti-ATV brain PK profiles in panels C and D. **(G)** Effective brain loss rate constants (kloss) estimated from the pulse–chase experiments (symbols; shaded regions indicate 90% confidence intervals) plotted as a function of receptor binding affinity (K_D_) and compared with model-predicted kloss values (lines) for ATV.TfR and ATV.CD98hc. Observed data in (A– D) are presented as mean ± SD (n = 4-5 animals/group/time point).

Simulations from the calibrated PK model were used to predict the plasma and brain PK profiles of ATV molecules in the absence of anti-ATV dosing (dashed traces in Fig. 4 A-D). Plasma PK measurements reflect the total huIgG concentrations, including free ATV or ATV–anti-ATV complexes in circulation. Following anti-ATV administration, plasma huIgG concentrations were higher than those predicted in the absence of anti-ATV intervention, consistent with the blockade of TfR- or CD98hc-mediated disposition (symbols in Fig. 4 A and B post anti-ATV administration). In parallel, brain huIgG concentrations declined more rapidly compared to the predicted traces without anti-ATV, reflecting the blockage of receptor-mediated brain uptake of ATVs following anti-ATV administration (Fig. 4 C and D). Notably, total rbIgG concentrations in the brain remained below ∼5 nM, consistent with limited brain penetration of anti-ATV antibodies primarily through non-specific uptake mechanisms (*SI Appendix,* Fig. S7).

To quantify the brain clearance of ATVs, we focused on the post anti-ATV phase of the brain PK profiles in Fig. 4 C and D. Since anti-ATV antibodies block receptor-mediated brain uptake of ATV molecules (as was confirmed earlier in Fig. 3 E and F), the time derivative of observed brain PK profile post anti-ATV provides a direct estimate of the brain clearance of the tested ATV variants (in units of concentration/time). To reduce noise sensitivity of numerical differentiation, we first fitted the post anti-ATV phase of each profile to a mono-exponential decay formula (solid traces in Fig. 4 C and D; *SI Appendix,* Table S7) and computed the time derivative from the fitted function (solid traces in Fig. 4 E and F).

### Receptor-specific brain clearance of ATV molecules

Analysis of the pulse–chase experiments revealed markedly higher brain clearance for ATV.TfR relative to ATV.CD98hc variants at comparable receptor binding affinities. For example, the estimated peak brain clearance flux for ATV.TfR (K_D_ = 100 nM, day 1) was >10x higher than that of ATV.CD98hc, despite a less than two-fold difference in receptor binding affinity (K_D_ = 170 nM, day 7; blue traces in Fig. 4 E and F). This was also reflected by the more rapid decline of ATV.TfR brain concentrations post anti-ATV administration, as opposed to the more sustained profiles for ATV.CD98hc variants (Fig. 4 C and D). These findings were also consistent with the brain clearance fluxes previously predicted by the PK model calibrated to the collective ATV dataset (Fig. 2 G and H). Furthermore, extending the PK model to incorporate anti-ATV administration (Materials and Methods; *SI Appendix,* Table S5), yielded simulated pulse–chase PK profiles that were in overall agreement with the experimental data, without requiring re-estimation of the previously calibrated ATV PK parameters (*SI Appendix,* Fig. S8). Importantly, since the pulse–chase data were not used for model calibration, these results further validate the predictive capability of the PK model calibrated to the collective ATV dataset to resolve the distinct brain fluxes of ATV molecules.

To further characterize the post–anti-ATV brain loss kinetics, the mono-exponential fits shown in Fig. 4 C and D were used to estimate an effective brain loss rate constant (termed *k_loss_*) for each ATV affinity variant (symbols in Fig. 4G; *SI Appendix,* Table S7). *k_loss_* represents the first-order brain loss rate constant and is inversely proportional to the apparent half-life of the molecules in the brain, enabling direct comparison of affinity-dependent brain loss kinetics independent of the initial brain concentration. ATV.TfR variants exhibited substantially higher *k_loss_* values (≥10x) than ATV.CD98hc (symbols). Additionally, despite overlapping confidence intervals, *k_loss_* estimates for ATV.TfR suggest an increasing trend with stronger receptor binding, consistent with enhanced TfR-mediated cellular internalization and degradation. In contrast, ATV.CD98hc variants exhibited uniformly low *k_loss_* values across the tested affinity range, suggesting limited brain loss and sustained brain retention.

Experimental *k_loss_* estimates were next compared with model-predicted values across a broader range of receptor binding affinities (Fig. 4G). Despite expected deviations at individual data points, the model recapitulated higher *k_loss_* of ATV.TfR relative to ATV.CD98hc while extending the analysis beyond the experimentally evaluated affinity range. At high K_D_ (weak receptor binding), model-predicted *k_loss_* values for both platforms converged toward a common baseline, consistent with minimal receptor-mediated transport. At low K_D_ (strong receptor binding), *k_loss_* predictions approached asymptotic limits, reflecting saturation of receptor-mediated brain loss for ATV.TfR and retention-associated mechanisms for ATV.CD98hc. Notably, although model predictions point to a modest increase in *k_loss_* with weaker CD98hc affinity, this trend was not fully resolved experimentally for the range of tested affinities. Interpretation of *k_loss_* estimates, particularly for the weak-affinity variants, should also be made with caution, as residual rbIgG concentrations in the brain become comparable to ATV concentrations and may bias the estimated values (*SI Appendix,* Fig. S7).

### Estimation of ATV brain uptake at the time of anti-ATV administration

Brain uptake can be estimated by combining the experimentally determined clearance flux with the net brain flux calculated from simulated brain PK profiles without anti-ATV administration (Fig. 5 A and B). Importantly, this approach is only directly applicable at the time of anti-ATV administration, when brain concentrations are comparable between animals receiving ATV alone and those receiving ATV followed by anti-ATV. Thereafter, inhibition of further brain uptake results in a smaller pool of residual ATVs available for clearance, such that the measured clearance flux may no longer represent the clearance that would have occurred in the absence of anti-ATV administration. By design, anti-ATV antibodies were administered near the predicted brain T_max_ of the corresponding ATV molecules to capture near-maximal brain clearance fluxes, while also taking advantage of the near-zero net flux at T_max_. Consequently, estimated brain uptake for ATV.TfR at day 1, and ATV.CD98hc at day 7 (Fig. 5 C and D) were similar to their previously estimated peak brain clearance (Fig. 4 E and F), indicating greater magnitude of brain uptake for ATV.TfR compared to ATV.CD98hc variants. However, these estimates correspond to different time points and, although expected to approximate peak brain clearance, may not capture peak brain uptake, particularly for ATV.CD98hc variants.

**Fig. 5.**
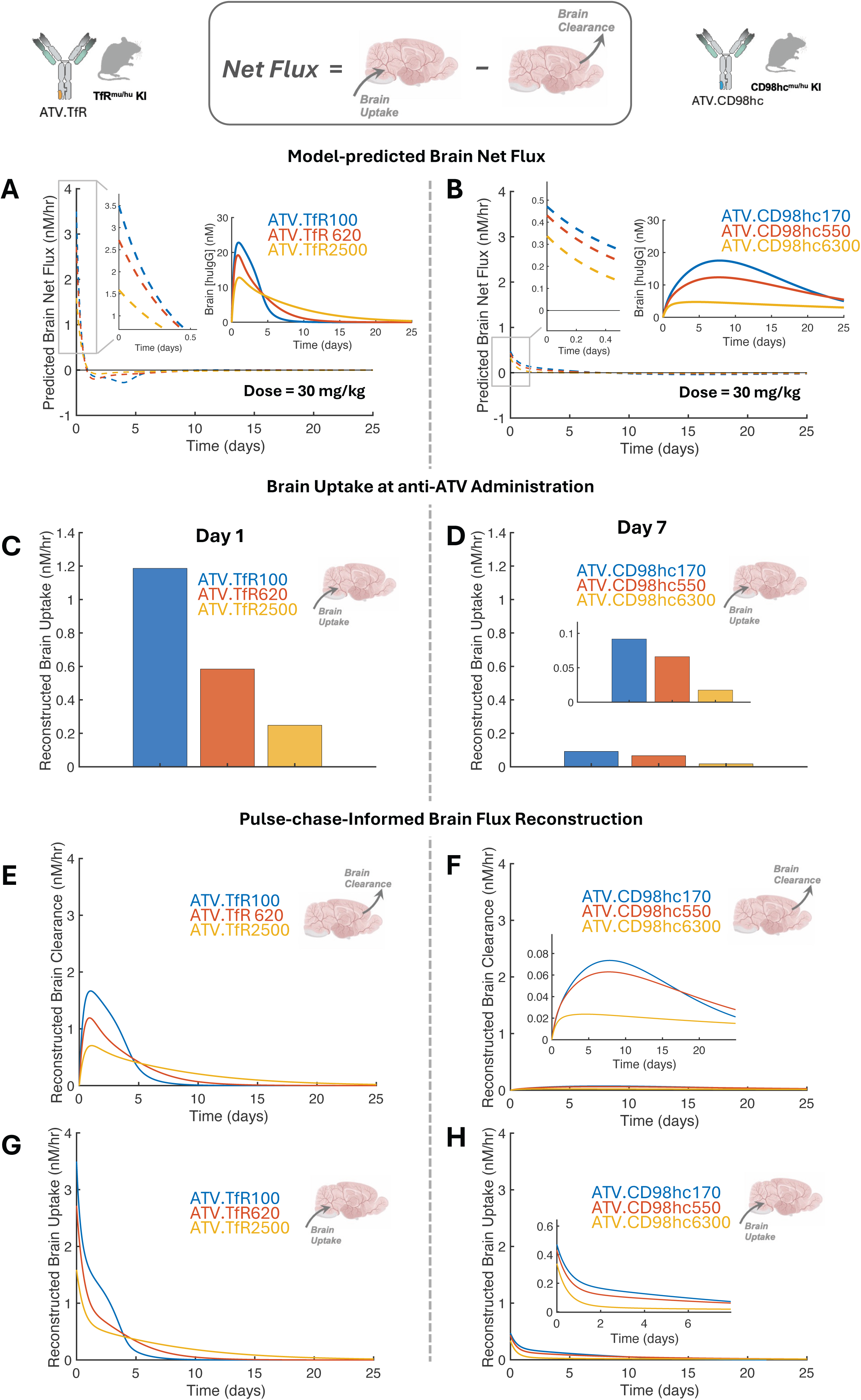
Pulse-chase-informed reconstruction of brain uptake and clearance fluxes for ATV.TfR and ATV.CD98hc over time. **(A,B)** Model-predicted net brain fluxes for ATV.TfR (A) and ATV.CD98hc (B) affinity variants, calculated from the corresponding simulated brain PK profiles (insets) in the absence of anti-ATV administration. **(C,D)** Brain uptake fluxes estimated at the time of anti-ATV administration (day 1 for ATV.TfR and day 7 for ATV.CD98hc) by combining experimentally estimated brain clearance fluxes (Fig. 4 E,F) with the corresponding model-predicted net brain fluxes (A,B). **(E,F)** Reconstructed time course of brain clearance fluxes of ATV.TfR (E) and ATV.CD98hc (F) variants informed by molecule-specific *k_loss_* estimates (Fig. 4G) **(G,H)** Reconstructed time course of brain uptake fluxes of ATV.TfR (G) and ATV.CD98hc (H) reconstructed by combining the estimated brain clearance fluxes (E,F) with the model-predicted net brain fluxes (A,B).

### Pulse-chase-informed reconstruction of brain fluxes of ATV molecules over time

Experimentally resolving brain uptake and clearance over the entire PK profile would require repeating the pulse-chase experiment at multiple intervention times and, at each time point, combining the estimated clearance flux with the corresponding net brain flux to reconstruct the full brain uptake profile. However, the molecule-specific effective brain loss rate constant (*k_loss_*) estimated from the pulse-chase experiments (Fig. 4G, *SI Appendix,* Table S7) provides an alternative means to reconstruct brain fluxes over time. Under first-order kinetics, *k_loss_* represents an intrinsic characteristic of brain loss for a given molecule, describing the fractional rate of clearance independently of the initial brain concentration. This allows brain clearance to be estimated at different time points without repeating the pulse-chase experiments. Assuming *k_loss_* remains constant over time, we reconstructed the time-dependent fluxes for each ATV variant in three steps: (i) estimate *k_loss_* from observed brain PK following anti-ATV administration (Fig. 4G); (ii) estimate brain clearance at each time point by applying the *k_loss_* to simulated brain concentrations without anti-ATV administration (i.e., brain clearance (t) ≃ *k_loss_* * brain [huIgG] (t); Fig. 5 E and F); and (iii) reconstruct brain uptake over time by combining the estimated brain clearance with the model-predicted net brain flux (Fig. 5 G and H).

Although the pulse-chase-informed reconstruction is subject to uncertainties in estimating *k_loss_*, its assumed time invariance, and potential differences between simulated and observed brain PK profiles, the results consistently indicated substantially greater brain uptake and clearance fluxes for ATV.TfR than ATV.CD98hc across the tested affinities. These findings were consistent with predictions from the mechanistic PK model (Fig. 1B), which explicitly accounts for nonspecific clearance, receptor engagement, and receptor-mediated degradation (Fig. 5 E–H and Fig. 2 E–H). This agreement further supports the use of the mechanistic model for subsequent flux analyses, particularly for dual-receptor-targeting ATVs.

### Flux-informed interpretation of dual ATV brain PK

Flux-based analysis of the brain uptake and clearance of single-receptor TfR- and CD98hc-targeting ATVs establishes a mechanistic framework to interpret the unique PK properties of dual ATVs. As was previously described (25), a representative dual ATV clone was administered intravenously at 25 mg/kg in TfR^mu/hu^/CD98hc^mu/hu^ double KI (DKI) mice and was compared to the corresponding single-receptor ATV formats (Fig. 6 A and B). Although the dual ATV exhibited lower systemic exposure than either ATV.TfR or ATV.CD98hc, consistent with increased peripheral engagement of both targets, brain concentrations were up to ∼3-fold higher than either single-receptor ATV and remained quantifiable throughout the 14-day study.

**Fig. 6.**
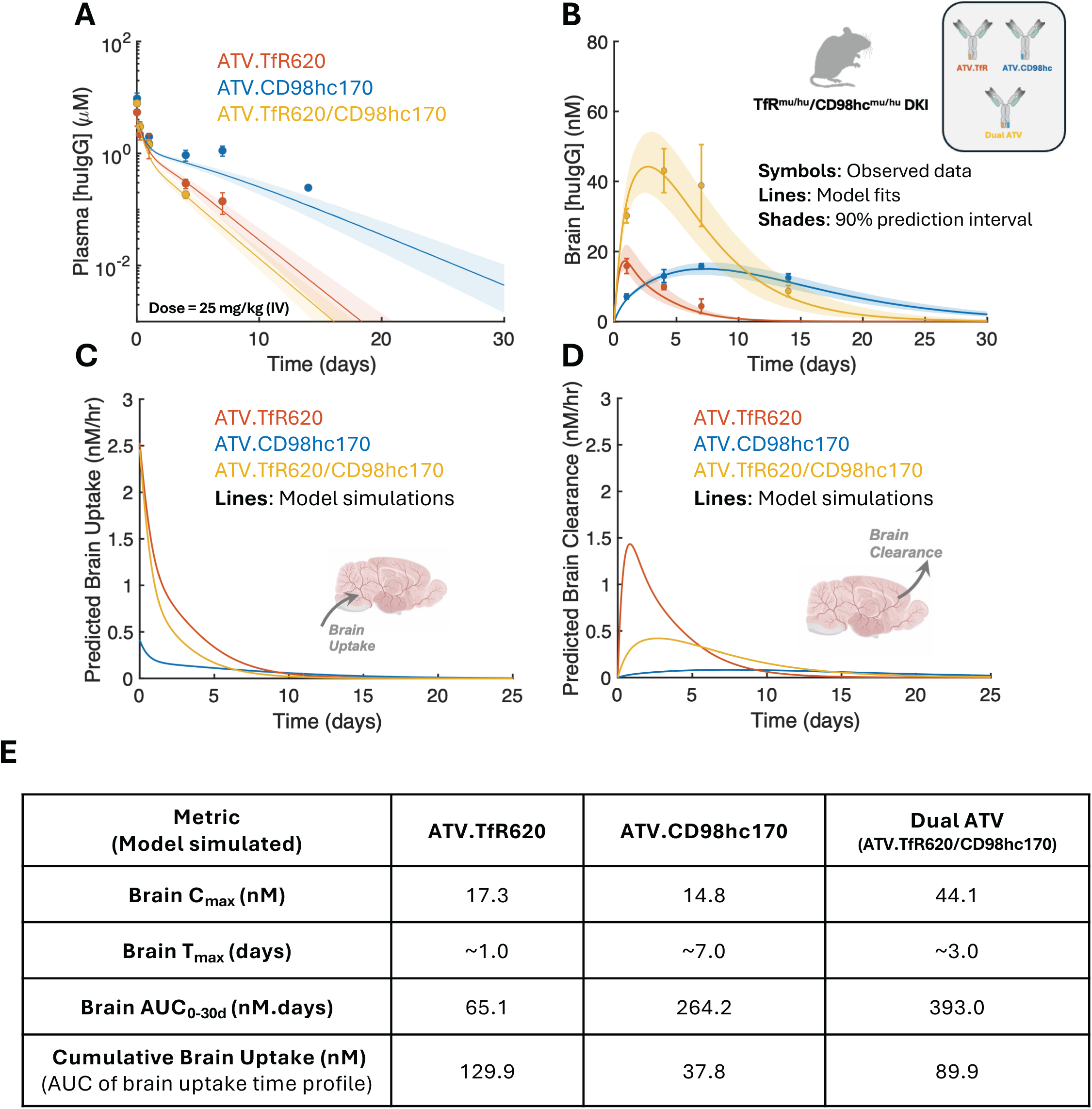
Model-informed flux analysis of a dual ATV construct compared with single-receptor ATV.TfR and ATV.CD98hc controls. **(A,B)** Observed plasma (A) and brain (B) PK profiles of the dual ATV, ATV.TfR, and ATV.CD98hc following a single 25 mg/kg intravenous dose in TfR^mu/hu^/CD98hc^mu/hu^ DKI mice. Symbols represent observed data (previously published by Wells et al. (25)), solid lines represent model fits, and shaded regions denote the 90% prediction intervals based on the uncertainty (%CV) in the estimated model parameters (*SI Appendix,* Table S4). Simulations of the dual ATV clone are done with the actual affinity values reported in Table S2. Observed data are presented as mean ± SD (n = 3-4 animals/group/time points). **(C,D)** Model-predicted brain uptake (C) and brain clearance (D) fluxes for the dual ATV and the corresponding single-receptor ATV.TfR and ATV.CD98hc controls. **(E)** Brain PK metrics for ATV.TfR, ATV.CD98hc, and the dual ATV, including brain C_max_, T_max_, AUC, and cumulative brain uptake (calculated as the area under the brain uptake flux–time profile), estimated from the corresponding model simulations.

We next used the calibrated PK model to determine whether the enhanced brain exposure of dual ATVs resulted from increased brain uptake, reduced brain clearance, or a combination of both (Fig. 6 C and D). As described in Materials and Methods, the dual ATV model assumes additive TfR- and CD98hc-mediated brain uptake, competitive receptor binding in the brain, and receptor occupancy governed by receptor expression and binding affinity. The model fits adequately captured the observed plasma and brain PK profiles of both single-receptor and dual ATV constructs (Fig. 6 A and B). Moreover, it reproduced the previously reported observed PK of dual ATVs across multiple receptor affinity combinations (25) and dose levels, including datasets reserved for model validation (*SI Appendix,* Figs. S1 and S2).

While dual-receptor engagement might be expected to enhance brain delivery by enabling parallel uptake pathways, modeling flux analysis suggests the possibility of an alternative mechanism. Simulations for the representative dual ATV clone predict a similar brain uptake for the representative dual ATV relative to the ATV.TfR control during the early phase after dosing, with uptake becoming comparable to or even lower than ATV.TfR at later time points (Fig. 6C). These results suggest that TfR-mediated transport remains the predominant contributor to brain uptake and that the higher systemic clearance of the dual ATV largely offsets any additional contribution from CD98hc-mediated uptake. In contrast, predicted brain clearance of the dual ATV was lower than ATV.TfR (Fig. 6D), indicating that CD98hc-mediated retention, via reduced catabolism, is the primary driver of sustained brain exposure. Together, these simulations suggest that the improved brain PK of dual ATVs arises predominantly from reduced brain clearance rather than enhanced brain uptake.

This distinction becomes particularly apparent when comparing cumulative brain uptake with conventional PK metrics such as brain C_max_ and AUC. Although the dual ATV achieved higher brain C_max_ and AUC than either single-receptor format, these metrics cannot distinguish whether increased exposure reflects greater drug delivery into the brain or prolonged retention. To address this, we calculated cumulative brain uptake as the integral of the brain uptake flux over time, representing the total amount of drug entering the brain over the evaluated period. Interestingly, despite its higher brain exposure, the dual ATV was predicted to have cumulative brain uptake comparable to, or slightly lower than, ATV.TfR (Fig. 6E), indicating that its increased brain AUC was primarily driven by reduced brain clearance rather than enhanced uptake.

### Predicted dual ATV brain fluxes across a wide range of receptor binding affinities

Simulating the brain fluxes across a broad matrix of TfR and CD98hc affinities reveals how tuning the receptor binding properties of the dual ATV can shape the balance between brain clearance and uptake (Fig. 7). While stronger binding to both receptors is predicted to increase brain C_max_ (Fig. 7A), the highest brain AUC values are predicted for combinations with strong CD98hc affinity and intermediate TfR affinity, indicating a trade-off between brain fluxes and systemic exposures (Fig. 7B). Interestingly, simulations predict that the maximum brain uptake flux of dual ATVs is governed predominantly by TfR targeting, with stronger TfR binding producing higher uptake and CD98hc affinity exerting only a modest influence (Fig. 7C). In contrast, brain clearance is lowest for dual ATV combinations with strongest CD98hc and weakest TfR binding, consistent with enhanced CD98hc-mediated retention and reduced TfR-mediated clearance (Fig. 7D). Notably, the cumulative brain uptake is predicted to be comparable or even lower for dual ATV pairings than for ATV.TfR alone (Fig. 7E), suggesting that the increased systemic clearance of dual ATVs offsets the modest gains in brain uptake achieved through dual receptor targeting. These results suggest that maximizing receptor engagement or peak brain concentration does not necessarily translate to greater cumulative delivery, underscoring the importance of tuning both uptake and retention mechanisms in the design of dual-receptor ATVs.

**Fig. 7.**
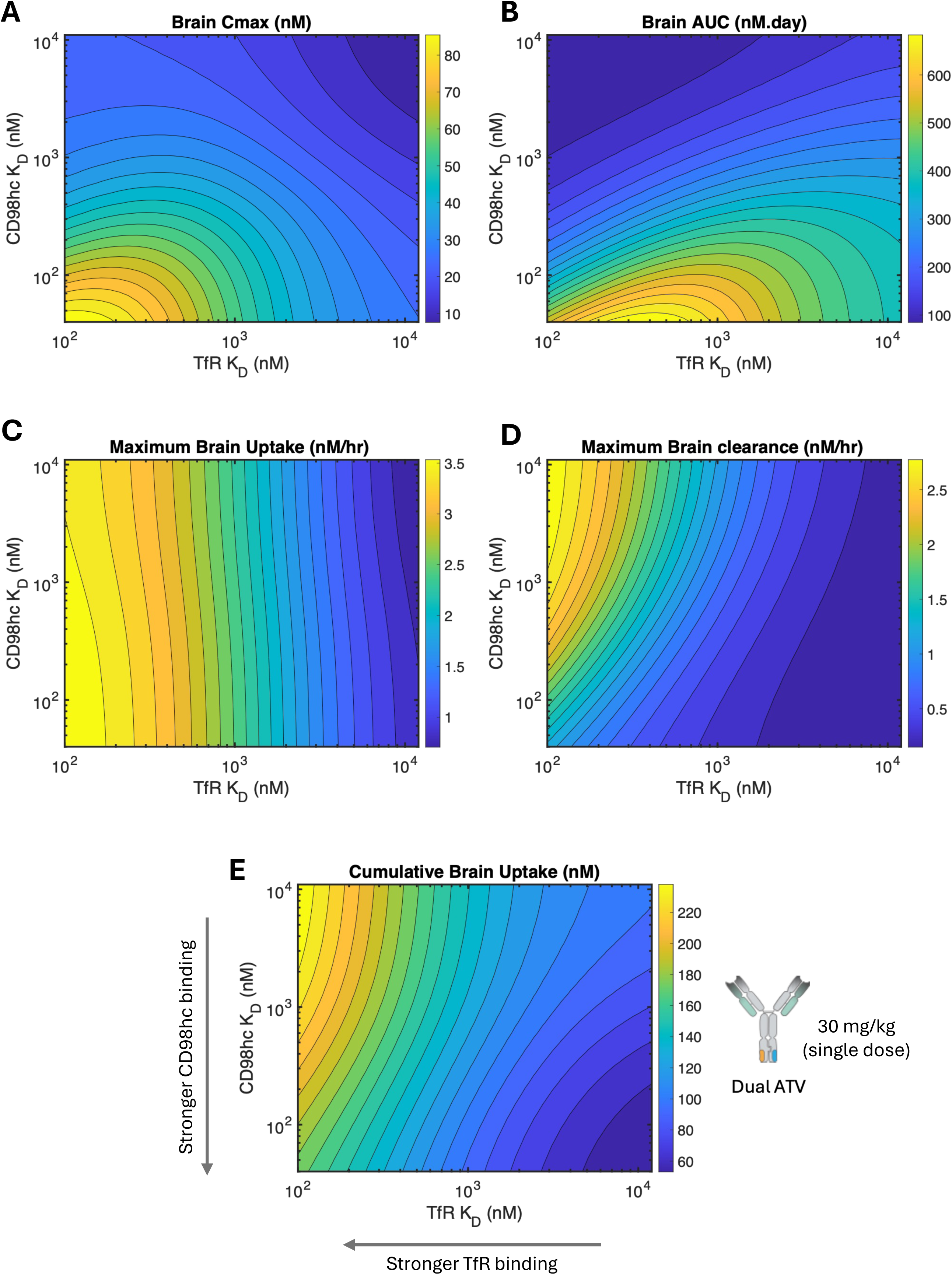
Model-predicted brain PK metrics and fluxes of dual ATVs across a matrix of TfR and CD98hc binding affinities. **(A–E)** Simulated relationships between TfR and CD98hc binding affinities (K_D_) and model-predicted brain PK metrics and fluxes for dual ATVs following a single 30 mg/kg intravenous dose in TfR^mu/hu^/CD98hc^mu/hu^ DKI mice. Heatmaps depict predicted **(A)** brain C_max_, **(B)** brain AUC, **(C)** maximum brain uptake flux, **(D)** maximum brain clearance flux, and **(E)** cumulative brain uptake (calculated as the area under the brain uptake flux–time profile).

## Discussion

Brain concentration is frequently used as a surrogate for CNS delivery; however, it reflects the net outcome of two dynamic processes: brain uptake and clearance. As we have shown in this study, similar brain concentrations at a given time point can arise from markedly different combinations of uptake and clearance, highlighting a limitation of conventional PK metrics that become especially consequential when comparing and further optimizing mechanistically distinct delivery platforms such as ATV.TfR and ATV.CD98hc. Resolving brain exposure into the underlying transport dynamics provides a mechanistic basis for the rational engineering of new brain-penetrant therapeutics. Beyond the brain, this flux-centric framework generalizes to any tissue in which competing transport processes govern exposure.

We mechanistically separated brain uptake and clearance by combining mathematical modeling with a targeted intervention. After initial systemic distribution, anti-ATV antibodies were used to block further receptor-mediated brain uptake, enabling direct measurement of brain clearance. When coupled with net brain flux, this allowed brain uptake to be estimated over time. The inverse strategy is also feasible: suppressing brain efflux, for example by targeting brain-retentive epitopes such as MOG, can enable direct measurement of uptake (48). Other complementary approaches have also emerged for probing flux. Veire *et al.* (2025) (46), for example, used Zr-89 and I-125 radiotracing to distinguish total brain entry from retention and degradation of TfR- and CD98hc-targeted antibodies, a critical dimension often missed by conventional PK studies. Despite using different constructs, their findings align with our observations that TfR-targeted molecules are associated with rapid brain clearance, whereas CD98hc contributes to parenchymal retention. Collectively, these studies highlight that brain concentration measurements alone can mask distinct and relevant transport dynamics, and that mechanistic flux analysis is needed to resolve pharmacologically meaningful transport behavior.

Dual TfR/CD98hc targeting ATVs further illustrate this principle. Although engaging two transport receptors might be expected to enhance brain delivery, our flux analysis suggests that dual targeting introduces uptake–clearance trade-offs that are not evident from the PK profiles. Specifically, the analysis predicted that dual TfR and CD98hc targeting does not enhance brain uptake compared to the single-receptor ATV.TfR construct, as the additional contribution of CD98hc-mediated uptake is largely offset by increased peripheral target-mediated clearance. Instead, analysis suggests the observed increase in brain exposure of dual ATVs primarily results from improved retention within the brain, mediated by CD98hc engagement. Although further experimental validation may be required for these conclusions, they underscore the need for a mechanistic framework to interpret dual- or multi-targeting outcomes. Determining whether receptor combinations act synergistically, neutrally, or antagonistically will require experimental and computational tools that resolve flux. Such tools can move the field beyond descriptive PK toward rational design, including tuning receptor affinity, defining when dual targeting helps or hinders delivery, and understanding how a therapeutic Fab affects uptake and clearance.

While this analysis provides mechanistic insight into brain uptake and clearance, it is limited by its reliance on brain homogenate data, which measures total brain concentration but not subregional, cellular, or compartment-specific distribution. The model therefore cannot resolve whether drug resides in vascular, interstitial, or intracellular spaces, even though these distinctions can strongly shape pharmacologic activity. Brain uptake is also modeled as the additive contribution of independent TfR- and CD98hc-mediated pathways, without accounting for potential nonadditive interactions, including avidity and cooperative or competitive receptor engagement. More detailed receptor-mediated transcytosis models could provide greater spatial resolution (51–54), but in practice are often limited by the availability and granularity of supporting data. Future extensions of this framework could incorporate capillary-depletion, immunohistochemistry, imaging, and pharmacodynamic (PD) data to better resolve brain biodistribution and link tissue exposure to target engagement and PD responses.

Importantly, total brain exposure alone does not define delivery success. Depending on the therapeutic application, biodistribution, exposure above a pharmacodynamic threshold, and compartmental localization may be equally or even more important for efficacy. In this regard, the modular TV platform offers a tunable design space in which uptake, clearance, and retention can be optimized to match the needs of a given therapeutic modality or target biology. More broadly, modular brain-targeted biologics using therapeutic Fabs, cargo domains, or multivalent targeting need to be evaluated individually, as each format can uniquely alter the balance between brain uptake and brain clearance. For example, an intracellularly retained cargo may benefit from rapid brain uptake and may tolerate high clearance, making it better suited to the TfR platform. Conversely, an extracellular antagonist may require longer parenchymal residence, making CD98hc TVs a better fit. Understanding these fluxes can help match molecular design to the desired PK profile.

Taken together, this work offers a generalizable framework for deconstructing brain exposure into its mechanistic components. By integrating mathematical modeling with targeted experiments aimed at interrogating the underlying brain fluxes, it becomes possible to design, compare, and optimize CNS delivery strategies with greater precision across molecular architectures and therapeutic targets.

## Materials and Methods

### Nomenclature

Throughout this manuscript, ATV molecules are referred to by simplified names based on receptor specificity and binding affinity (*SI Appendix,* Table S1). For example, ATV.TfR620 denotes a TfR-targeting ATV with a measured monovalent TfR binding affinity of 620 nM. Similarly, ATV.CD98hc170 denotes a CD98hc-targeting ATV with a measured monovalent CD98hc binding affinity of 170 nM. Dual ATVs are designated using both receptor specificities (e.g., ATV.TfR620/CD98hc170), where the affinity values correspond to the “expected” binding affinities of the respective TfR- and CD98hc-targeting TVs, derived from the corresponding single-receptor control ATVs. Experimentally measured (“actual”) binding affinities of the dual ATVs are provided in *SI Appendix,* Table S2, which may differ modestly from the expected values. Molecular complexes are denoted using a hyphen (e.g., ATV–anti-ATV).

### Animal Care

All procedures in animals were performed under ethical regulations and protocols approved by Denali Therapeutics Institutional Animal Care and Use Committee. Housing maintained a 12-h light/dark cycle and had access to water and chow (LabDiet 5LG4, Irradiated) *ad libitum*. Temperature and humidity in all animal rooms were monitored daily by Thermo Scientific™ InSight. The temperature range was 18.3–23.3 °C and the humidity range was 30–70%.

### Dosing, blood and tissue collections

All mouse *in vivo* studies utilized male and female mice with C57BL/6J background (The Jackson Laboratory). Sex and age were distributed equally across all *in vivo* studies. Administration was done via tail vein injection (IV) and in-life sampling was done via submandibular or submental bleeds. At terminal time points, animals were deeply anesthetized using 2.5% Avertin via IP and collection of blood was done via cardiac punctures. Blood was collected in EDTA tubes and spun immediately at 12,700 x g for 7 minutes to separate plasma which was used for quantification of huIgG. Animals were then transcardially perfused with PBS and tissues were collected and flash frozen. Frozen tissues were homogenized for 6 min at 30 Hz using a Qiagen TissueLyser with 5 mm steel beads in 10x tissue weight of lysis buffer containing 1% NP40 in PBS with protease inhibitors. Specifics of sample sizes, dosing, collection times, and model fitting is reported in the figure legends.

### Quantification of huIgG

Antibody concentrations were quantified using a polyclonal anti-human IgG sandwich ELISA assay. ELISA plates (384-well, Thermo #464718) were coated overnight at 4°C with donkey anti-human IgG (JIR #709-006-098) diluted to 1 ug/mL with sodium bicarbonate solution (Sigma #C3041-50CAP). Plates were then blocked with blocking buffer (PBS + 0.05% Tween 20 + 5% Sigma BSA) for 2 hr at room temperature with shaking (500 rpm). Plates were then washed three times using wash buffer (PBS + 0.05% Tween 20). Standard and samples were diluted in PBS + 0.05% Tween 20 and 1% BSA. The standard curve range for the assay was 0.41 - 1500 ng/mL or 0.003 - 10 nM (BQL < 0.03 nM for plasma and BQL < 0.3 nM for tissues). Standards and samples were incubated at room temperature for 2 hrs with shaking (500 rpm). Plates were then washed three times with wash buffer. A goat anti-human IgG antibody (JIR #109-036-098) was diluted with PBS + 0.05% Tween 20 and 1% BSA at 1:40,000 and incubated for 1 hr at room temperature with shaking (500 rpm) for detection. The plates were then washed three times, developed using a TMB solution (Thermo #34028) over a 5 – 10 min incubation, quenched using 4N H_2_SO_4_ (Life Tech #SS04) and absorbance at 450 nm was then measured in a Molecular Devices SpectraMax 384 Plus plate reader.

### Quantification of dual ATV in tissues

Total test article concentrations in brain lysate samples were quantified using an anti-human IgG sandwich electrochemiluminescence immunoassay (ECLIA) on the Meso Scale Discovery (MSD) platform. All assay steps were performed at ambient temperature with shaking, as appropriate. Briefly, an MSD GOLD 96-well small-spot streptavidin-coated microtiter plate (Meso Scale Discovery, L45SA) was blocked with 1% casein-based PBS blocking buffer (Thermo Scientific, 37528) for approximately 1 hour. After blocking and washing, biotinylated goat anti-human IgG (SouthernBiotech, 2049-08) was added at a working concentration of 0.5 µg/mL to coat the plate, followed by a 1–2 hour incubation.

Test samples, diluted at a minimum required dilution (MRD) of 1:100 in 0.5% casein-based PBS assay buffer, were then added to the plate and incubated for 1–2 hours to allow capture of the analyte. A pre-adsorbed, ruthenylated (SULFO-TAG) goat anti-human IgG detection antibody (Meso Scale Discovery, R32AJ) at 0.5 µg/mL was subsequently applied and incubated for approximately 1 hour. The ECL signal was generated using 1X MSD Read Buffer T (R92TC) and measured as electrochemiluminescence units (ECLU).

Brain lysate samples required an MRD of 50 and demonstrated a dynamic range of 4.9 – 2500 ng/mL, based on a 10-point, 1:2 serially diluted standard curve plus a blank matrix control. Sample concentrations were calculated from the assay-specific calibration curve using a weighted four-parameter logistic (4-PL) regression model. The back-calculated concentrations (ng/mL) were then converted to nanomolar (nM) for final reporting.

### Quantification of anti-ATV Rabbit IgG (rbIgG)

The total test article concentrations in brain lysate samples were quantified using an anti-rabbit IgG sandwich-format electrochemiluminescence immunoassay (ECLIA) on a Meso Scale Discovery (MSD) platform. Briefly, 1% casein-based PBS blocking buffer (Thermo Scientific, Reference No. 37528) was added to an MSD GOLD 96-well small-spot streptavidin-coated microtiter plate (Meso Scale Discovery, Reference No. L45SA) and incubated for approximately 1 hour. Following the plate blocking and wash steps, biotinylated goat anti-rabbit IgG (SouthernBiotech, Reference No. 4030- 08) at a working concentration of 1.0 μg/mL was added to coat the assay plate and allowed to incubate for 1-2 hours. Subsequently, test samples, which were pre-diluted accordingly to fit into the assay standard curve range, were added to the assay plate and allowed to incubate. Following the 1-2 hours incubation in the capture step, a pre-adsorbed secondary ruthenylated (SULFO-TAG) goat anti-rabbit IgG antibody (Meso Scale Discovery, Reference No. R32AB) at a working solution of 0.5 μg/mL was added to the assay plate and incubated for approximately 1 hour. An assay read buffer (1X MSD Read Buffer T, R92TC) was then added to generate the electrochemiluminescence (ECL) assay signal, expressed in ECL units (ECLU). All assay reaction steps were performed at ambient temperature and with shaking on a plate shaker (where appropriate). The standard curve range for the assay was 1.22 – 1250 ng/mL in a neat plasma matrix and 0.24 – 250 ng/mL in a neat brain lysate matrix. Concentrations were back-calculated off the assay-specific calibration standard curve, which was fitted with a weighted (1/y^2^) four-parameter logistic regression using Microsoft Excel. The sample back-calculated concentrations in ng/mL were subsequently converted to nanomolar (nM) as the final sample results.

### Noncompartmental Analysis

Non-compartmental analysis was performed using Dotmatics software 5.5 (Boston, Mass) using a non-compartmental approach consistent with the intravenous route of administration. Parameters were estimated using nominal sampling times relative to the start of each administration. Samples that were below the quantitation limit (BQL) were omitted. The model linear up log down was used to calculate AUC exposures.

### Expression and purification of CD98hc and TfR apical domain

DNA encoding the extracellular domain (ECD) of human CD98hc (UniProt Q01650) was cloned into a pRK5 mammalian expression vector downstream of a mouse κ light chain signal peptide. A TEV- cleavable Avi-tag and hexahistidine tag (Avidity, LLC) was added in-frame at the C-terminus for purification and detection. TfR constructs were generated similarly, with the sequence for a circularly permutated human TfR apical domain (4) inserted into the same expression cassette.

### Expression and purification of ATVs and antibodies

Sequences encoding the heavy and light chains of antibodies and ATV variants were cloned into a pRK5 mammalian expression vector, using anti-DNP Fab sequences previously described (24). LALAPG mutations were incorporated into heavy chain sequences to attenuate FcγR binding and knob (T366W) with hole (T366S/L368A/Y407V) (4) mutations were incorporated into ATV variants to facilitate heterodimerization and achieve monovalent RMT binding or heterobivalent TfR/CD98hc binding on a single ATV. Resulting plasmids were used to transfect Expi293FTM cells (Gibco/Thermo Fisher A14527) and/or CHOK1 glutamine synthase-knock out cell line (Horizon) using standard methods. Recombinant antibodies and ATVs were purified from culture supernatant using protein A affinity chromatography (mAb Select SuRe, Cytiva), polished by size exclusion chromatography (Superdex 200, Cytiva), and formulated in 1xPBS or 10 mM sodium acetate with 6% sucrose, pH 5.5. Purity of the final molecules was confirmed to be >95% by HPLC-SEC and SDS-PAGE, and identities were confirmed by LC/MS. Proteins were stored at 4 °C or flash frozen and stored at -80 °C until use. Material purity and affinity were confirmed using SEC, LC/MS, endotoxin testing (<1EU/mg using Charles River Endosafe MCS instrument) and SPR prior to animal dosing.

### Animal immunizations with ATV.TfR and ATV.CD98hc

Balb/c and CD1 mice (Charles River) were immunized with 20 µg ATV.CD98hc or 20 µg ATV.TfR formulated with 10 µg c-di-GMP Vaccigrade (Invivogen, vac-nacdg emulsified in Sigma Adjuvant System (Sigma Aldrich, S6322). Mice were boosted multiple times over four weeks with 20 µg of the respective immunogen in the same adjuvant formulation.

### Hybridoma antibody discovery

For anti-ATV.TfR antibody, antibody discovery was performed at BBI-Solutions Maine (formerly Maine Biotechnology Services). For anti-ATV.CD98hc, lymph nodes and splenic B cells were isolated by negative selection and fused with SP2/0 cells by electrofusion to generate hybridomas (55). The target antigen was ATV.CD98hc, with the 170 nM sequence representing the entire TV family. Hybridomas were selected in HAT medium (Sigma Aldrich, 0262), screened for antibody secretion specific to ATV.CD98hc-Fc and counter screened against human Fc controls to ensure TV patch specificity. Supernatants were subcloned into full-length mouse IgG2a and further characterized by ELISA. Assays were performed in flat-bottom 96-well plates (Nunc Maxisorp, Affymetrix) coated overnight at 4 °C with 2 µg/mL antigen in PBS. Plates were washed three times with PBS, 0.05% Tween-20 (PBS-T, (pH 7.4), blocked for 1 h at room temperature with 3% BSA in PBS-T, and washed four times. For screening, 25 µL hybridoma supernatant from culture wells was added to antigen-coated plates and incubated for 1 h at room temperature with gentle agitation. Plates were washed three times, incubated for 1 h with goat anti-mouse HRP-conjugated antibody (Jackson ImmunoResearch, 115-035-164), washed again, and developed with TMB substrate (Surmodics, TMBW-1000-01). Reactions were stopped with TMB stop solution (Surmodics, LSTP-1000), and absorbance was measured at 450 nm.

### Phage display library construction for anti-ATV.CD98hc discovery

RNA from spleens and lymphocytes of ATV.CD98hc-immunized animals was isolated using QIAshredder columns (Qiagen, 79656) and the PureLink RNA Mini Kit (Invitrogen, 12183020). cDNA was synthesized from random hexamers with the SuperScript IV kit (Invitrogen, 18091050). Variable heavy (VH) and light (VL) chain sequences were amplified using mouse-specific primers(56). VL and VH regions were assembled by overlap extension PCR to generate scFv fragments (VL-linker-VH) flanked by SfiI sites. PCR products were cloned into an ampicillin-resistant phagemid vector via SfiI digestion, encoding scFvs with C-terminal 6×His and c-Myc tags fused to the M13 phage pIII protein. The library was transformed into TG1 electrocompetent cells (Biosearch Technologies, 60502-1), yielding ∼1 × 10¹⁰ transformants.

### Phage display for anti-ATV.CD98hc discovery

*E. coli* carrying the library were cultured in TB medium (1% glucose, 200 μg/mL carbenicillin) at 37 °C to OD600 = 0.5, then infected with 1 × 10¹¹ pfu/mL M13K07 helper phage (NEB, N0315S) and induced with 1 mM IPTG. After 2 h at room temperature with shaking, 12.5 μg/mL kanamycin was added, and cultures were grown overnight for phage production. Phage were precipitated with 20% PEG-8000/2.5 M NaCl, tittered on TG1, and used for panning. This amplification panning cycle was repeated for four rounds. In Round 1, 2 μg/mL anti-c-myc antibody (Invitrogen) was coated on high-binding plates to deplete phage expressing truncated or absent scFvs. Rounds 2–4 involved solution-phase binding of phage to biotinylated ATV.CD98hc-Fc captured on streptavidin magnetic beads (Thermo Scientific, 88816). Beads (100 μL) were blocked with 2% BSA in PBS-T (PBS, 0.05% Tween-20), and phage were preincubated with blocked beads for 30 min. Fc-binders were removed by depletion with Fc- coated beads (10 μg/mL), blocked and washed as above, for 1 h at room temperature. Unbound phage were then incubated with biotinylated ATV.CD98hc-Fc (1 mL final volume), followed by capture on beads (<5 min). Beads were washed 3–4 times with PBS-T, and bound phage were eluted with 0.5 mL of 10 mM glycine (pH 2.0, 10 min), then neutralized with 50 μL 1 M Tris-HCl. Across Rounds 2–4, antigen concentrations were decreased (to 10, 5, and 1 μg/mL). In parallel, eluates were also recovered with 0.1 M HCl (pH 2.2), neutralized 1:10 with 1 M Tris base, and amplified in TG1. Random clones from Rounds 3 and 4 were sequenced, and unique scFvs were evaluated by monoclonal phage ELISA. Plates were coated overnight at 4 °C with 1 μg/mL antigen (ATV.CD98hc-Fc variants, anti-c-myc, or Fc), washed with PBS-T, and blocked with PBS-T/2% BSA. Phage were incubated 1 h at room temperature, washed, and detected with anti-M13 pVIII- HRP (GE Healthcare, 1:2000 in PBS-T/2% BSA). After washing, plates were developed with TMB substrate (Thermo Scientific), quenched with 1 N HCl, and absorbance was measured at 450 nm.

### Cell binding and competition on cells

Cell binding and competition were evaluated by immunocytochemistry with high-content imaging. HEK293T cells (ATCC, CRL-3216), which endogenously express TfR and CD98hc, or CHO parental cells (ChemPartner, discontinued), were seeded at 30,000 (HEK293T) or 15,000 (CHO) cells/well in 96-well poly-D-lysine plates (Fisher Scientific/PerkinElmer, 6055302) and incubated overnight at 37 °C. Cells were treated for 1 h at 37 °C with 250 nM anti-ATV.TfR, anti-ATV.CD98hc, anti-ATV.TfR-ATV.TfR, or anti-ATV.CD98hc-ATV.CD98hc, then fixed with 4% paraformaldehyde and blocked in PBS with 5% BSA and 0.3% Triton.

Cells were stained with AlexaFluor®488-conjugated anti-human IgG (1:1000, Jackson ImmunoResearch, 109-545-003), DAPI, and Deep Red Cell Mask (ThermoFisher, C10046). For each replicate, ≥ 20 fields of view were acquired at 40 × magnification on the Opera Phenix High-Content Imaging System (PerkinElmer). Images were analyzed using Harmony software (PerkinElmer v4.9).

### Crystal structure of ATV.TfR–anti-ATV.TfR complex

The ATV.TfR100 (Fc only)–anti-ATV.TfR Fab complex was assembled by mixing ATV.TfR100 (Fc only) with an excess of anti-ATV.TfR Fab and purified by SEC. The complex was crystallized at a concentration of 11 mg/mL in 20% PEG 3350 by sitting well drop vapor diffusion at 18°C. Crystals appeared after 4 days and were cryoprotected in mother liquor supplemented with 20% ethylene glycol. Diffraction data were collected at the Canadian Light Source beamline CMCF-ID. Data were processed to a resolution of 2.0 Å in DIALS 2.8, and phases were obtained through molecular replacement with Phaser 2.8.3, using the CH2 and CH3 domains of TV35 (PDB: 6W3H) (4), as well as the variable and constant domains from an internal Fab crystal structure as search models. Model building and refinement were performed using COOT 0.9.8.1, ISOLDE 1.7, and REFMAC5 (*SI Appendix,* Table S8).

### Crystal structure of ATV.CD98hc–anti-ATV.CD98hc complex

The ATV.CD98hc170 (Fc only)–anti-ATV.CD98hc Fab complex was assembled by mixing ATV.CD98hc170 (Fc only) with an excess of anti-ATV.CD98hc Fab and purified by SEC. The complex was crystallized at a concentration of 26.5 mg/mL in 29% PEG 1500 by hanging drop vapor diffusion at 18 °C. Crystals appeared after 5 days and were cryoprotected in mother liquor supplemented with 20% ethylene glycol. Diffraction data were collected at the Canadian Light Source beamline CMCF-BM. Data were processed to a resolution of 2.7 Å in DIALS 2.8, and phases were obtained through molecular replacement with Phaser 2.8.3, using the CH2 and CH3 domains of ATV.CD98hc (PDB: 8G0M), as well as the variable and constant domains from an internal Fab crystal structure as search models. Model building and refinement were performed using COOT 0.9.8.1, ISOLDE 1.7, and REFMAC5 (*SI Appendix,* Table S8).

### Affinity determination by KinExA (Kinetic Exclusion Assay)

Kinetic exclusion affinity analysis measures both equilibrium binding and association kinetics without perturbing solution equilibrium, providing a physiologically relevant view of binding interactions. For anti-ATV.CD98hc and anti-ATV.TfR antibodies, Fabs were captured on azlactone beads and equilibrated with Fc-only ATV.CD98hc (8 nM -200 nM) or ATV.TfR family members (250 pM - 1 nM). Binding was detected using anti-human Fcγ-specific IgG (Jackson ImmunoResearch, 109-605-008). Fluorescent signals were converted to voltage, directly proportional to the concentration of free binding partner. All experiments were performed at Sapidyne Instruments.

### Affinity determination by surface plasmon resonance: I) ATV.CD98hc

Affinities of ATVs for human CD98hc were determined by surface plasmon resonance (SPR) using a BiacoreTM 8K instrument (Cytiva). ATV variants were immobilized on a Series S CM5 sensor chip (Cytiva, 29149603) using a human Fab capture kit (Cytiva, 28958325) at a concentration of 10 μg/mL and a flow rate of 10 μL/minute for one minute. 3-fold serial dilutions of recombinant human CD98hc extracellular domain (24) from 6 μM to 24.5 nM were injected at a flow rate of 30 μL/minute for five minutes followed by a 10-minute dissociation in 1X HBS-EP+ running buffer (Cytiva, BR100826). Kinetic analysis was performed using a 1:1 Langmuir kinetic binding model to evaluate k_on_, k_off_, and K_D_ for interactions stronger than 600 nM. A steady state model analysis was used to determine K_D_ values for weaker interactions, using the average of 3 independent experiments. Biacore Insight Evaluation software (version 2.0.15.12933) was used to analyze the data.

### Affinity determination by surface plasmon resonance: II) ATV.TfR apical

The affinity of ATV:TfR variants for recombinant TfR apical domain was determined by surface plasmon resonance using a Biacore^TM^ 8K instrument (Cytiva). Anti-human Fab (human Fab capture kit from GE Healthcare) was immobilized on Biacore™ Series S CM5 sensor chips. ATV (5 µg/mL) was captured for 1 minute on each flow cell, and serial 3-fold dilutions of hTfR apical domain (4) were injected at a flow rate of 30 µL/min. Each sample was analyzed with a 3-minute association and a 3-minute dissociation. After each injection, the chip was regenerated using 10 mM glycine-HCl (pH 2.1). Binding response was corrected by subtracting the RU from a flow cell capturing an irrelevant IgG at similar density. A steady state model analysis was used to determine K_D_ values for weaker interactions, using the average of 3 independent experiments. Biacore Insight Evaluation software (version 2.0.15.12933) was used to analyze the data.

### Mathematical modeling

A schematic of the ATV PK model (including ATV.TfR, ATV.CD98hc, and dual ATV constructs) is shown in Fig. 1B. Systemic PK was represented by a two-compartment model incorporating distribution into peripheral tissues, linear non-specific clearance, and receptor-mediated disposition through binding to TfR and CD98hc receptors. Dual ATV molecules were assumed to bind independently to each receptor, such that only one receptor could be engaged at a time, with no secondary or ternary complex formation (25). Additionally, receptor interactions were modeled as additive, without cooperativity or avidity effects.

Receptor-mediated clearance from systemic circulation was modeled under the quasi–steady-state approximation (57), with TfR- and CD98hc-mediated elimination each described by an independent Michaelis–Menten kinetics. For receptor *i* (where *i* ∊ [TfR, CD98hc]), V_max_ denotes the maximal clearance capacity via receptor *i*, with the Michaelis-Menten constant, K_m,i_, representing the apparent affinity governing receptor-mediated elimination of the ATV molecule, parameterized as a function of the receptor binding affinity (*α_i_*_,*c*_*K_D_*_,*i*_; *SI Appendix,* Table S4).

Brain uptake was modeled as the sum of (i) a non-specific uptake, representing passive entry processes across the blood–CSF barrier, and (ii) receptor-mediated transcytosis across brain endothelial cells at the BBB. TfR– and CD98hc–mediated transcytosis was represented by independent Michaelis–Menten terms, with additive contributions from each receptor pathway with no assumed avidity effects. Additionally, to account for differences in underlying biological processes governing receptor-mediated systemic clearance and transcytosis across the BBB, separate apparent affinity parameters (Km) were estimated for each process while maintaining their dependence on the corresponding receptor binding affinity (*α_i_*_,*c*_*K_D_*_,*i*_ vs. *α_i_*_,*b*_*K_D_*_,*i*_; *SI Appendix,* Table S4).

Within the brain compartment, ATV kinetics were modeled using a full TMDD framework (57) with explicit representation of binding ATV molecules to the TfR and CD98hc receptors. Elimination from the brain was modeled by (i) first-order non-specific loss of free ATV and (ii) degradation of ATV–receptor complexes.

### Model calibration and parameter estimation

The PK model was developed using the SimBiology® Toolbox (MATLAB R2024b, MathWorks). Calibration of model parameters was performed against PK data from control huIgG and ATV molecules in preclinical studies in TfR^mu/hu^ KI, CD98hc^mu/hu^ KI, and TfR^mu/hu^/CD98hc^mu/hu^ DKI mouse models. To evaluate the predictive performance of the model, a subset of the dual ATV PK data was withheld as a validation dataset and excluded from calibration (*SI Appendix,* Table S1).

Parameter estimation was conducted using the global optimization algorithm *patternsearch* in a stepwise procedure, beginning with plasma PK and subsequently incorporating brain PK. A pooled analysis was utilized to estimate a single parameter set capturing the totality of the calibration dataset. For control huIgG, receptor binding processes were assumed to be absent. For systemic PK parameters, data from both ATV and control huIgG molecules were fit together. Plasma PK data were first log10-transformed and were fitted using the constant error function. For brain PK calibration, non-specific brain uptake and loss parameters were first estimated using the PK data from control huIgG and were then fixed during calibration of ATV molecules. Combined error function was used for estimation of brain PK parameters of ATV molecules.

Model fits to the observed data and goodness-of-fit (GOF) metrics showed reasonable agreement between simulations and PK data across both calibration and validation datasets (*SI Appendix,* Figs. S1-3). Some discrepancies with individual data points were observed, which is expected given the pooled analysis as well as inter-study variability. In addition, the relatively narrow confidence intervals around the mean of the estimated parameters, reflected by tight coefficients of variation (CV, 95%), indicate that the model achieved a satisfactory fit across studies. For each simulation scenario, the 90% prediction bands reflecting parameter uncertainty were generated by simulating 300 plasma and brain PK parameter sets sampled within their %CV-derived confidence intervals using Latin Hypercube Sampling. A list of model state variables and parameters is provided in *SI Appendix,* Tables S3 and S4.

### Modeling ATV–anti-ATV interaction

The systemic PK of anti-ATV antibodies was represented by a two-compartment model with linear clearance and distribution into peripheral tissues. Brain uptake and loss of anti-ATVs were represented by non-specific uptake and first-order loss mechanisms, respectively. Because anti-ATV.TfR and anti-ATV.CD98hc do not bind either TfR or CD98hc, receptor-mediated processes were not included. Furthermore, the comparable plasma PK profiles of anti-ATV.TfR and anti-ATV.CD98hc (*SI Appendix,* Fig. S6G), which were also consistent with that of anti-DNP (Fig. 2A), supported fixing the general systemic disposition (*V_c_*, *V_p_*, *CL*, and *CL_D_*) and the non-specific brain uptake and loss parameters (*k_up_*_,*ns*,*b*_, *k_loss_*_,*ns*,*b*_) to those of the ATV model, thereby avoiding further model parameterization (*SI Appendix,* Table S5). Reversible binding between anti-ATV antibodies and ATVs was modeled using the mean of the corresponding association and dissociation rate constants reported in *SI Appendix,* Table S6. ATV–anti-ATV complexes formed in the central, peripheral, or brain compartments were assumed to follow the same disposition as free anti-ATV antibodies. Brain uptake of ATV–anti-ATV complexes from systemic circulation was assumed to be neglibible.

### Model equations

The full system of ordinary differential equations (ODEs) governing the ATV PK model are presented below. Equations are written in their general form for a dual ATV construct, in which both TfR- and CD98hc-mediated transport pathways are present. Receptor-specific terms are activated using binary switch variables (*δ*_TfR_ and *δ*_CD98hc_), such that only the corresponding transport processes are active for single-receptor ATV.TfR or ATV.CD98hc molecules, whereas both pathways are active for the dual ATV. Similarly, anti-ATV binding reactions are written using a generic notation, with the corresponding association and dissociation rate constants (*k*_on,antiATV_ and *k*_off,antiATV_) representing either anti-ATV.TfR or anti-ATV.CD98hc, depending on the simulation being performed.

### Central compartment (systemic circulation)

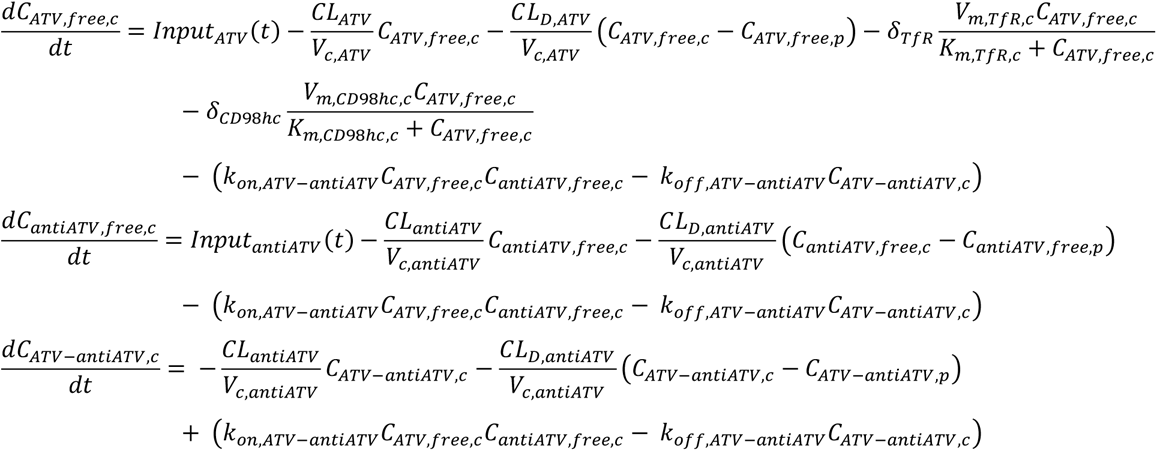

### Peripheral compartment

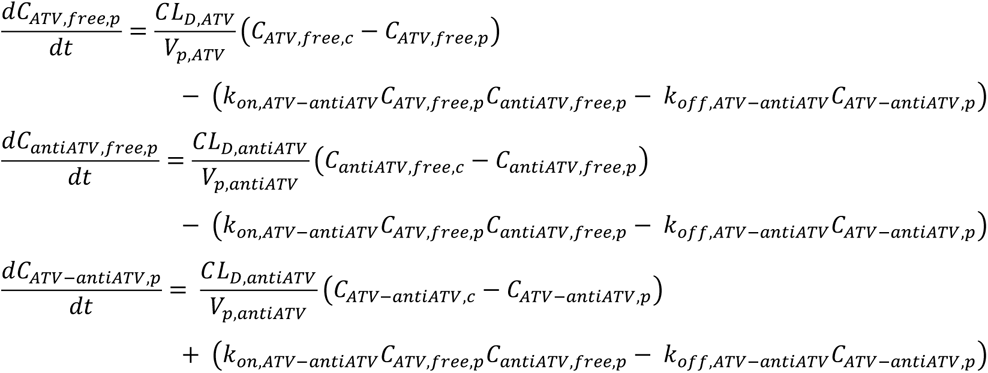

### Brain compartment

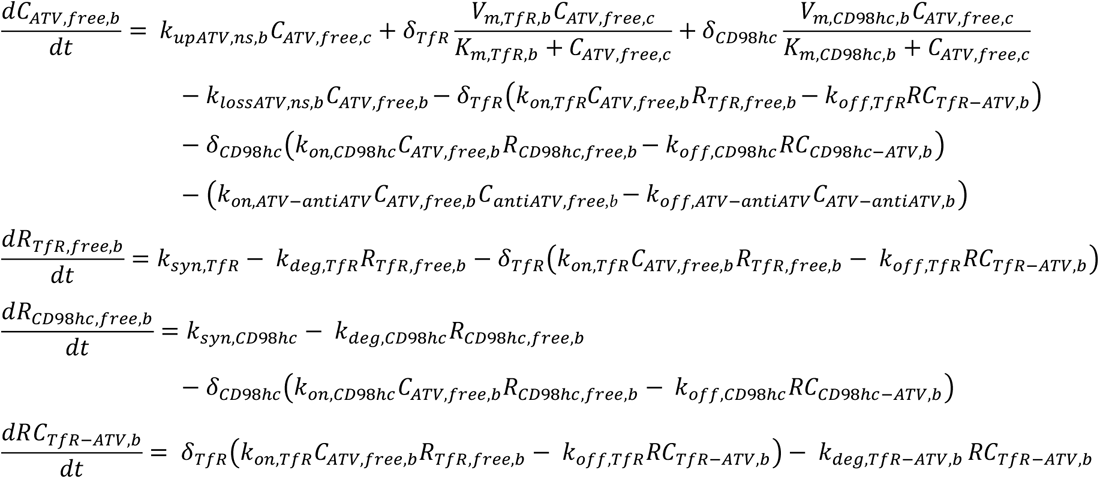

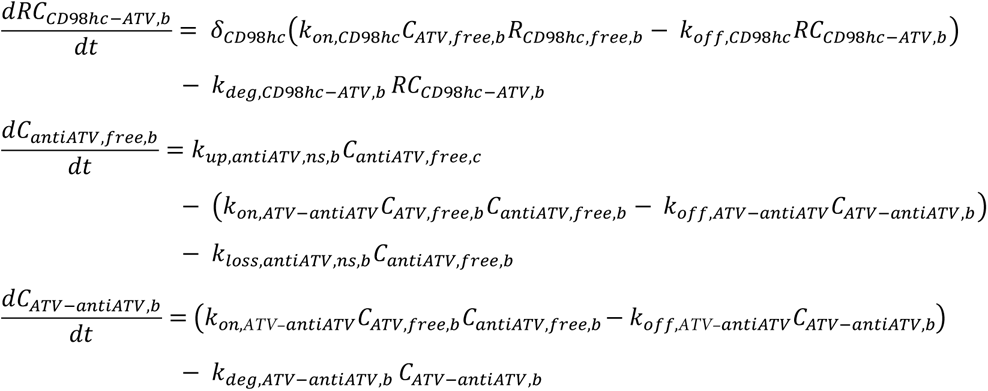

Where,

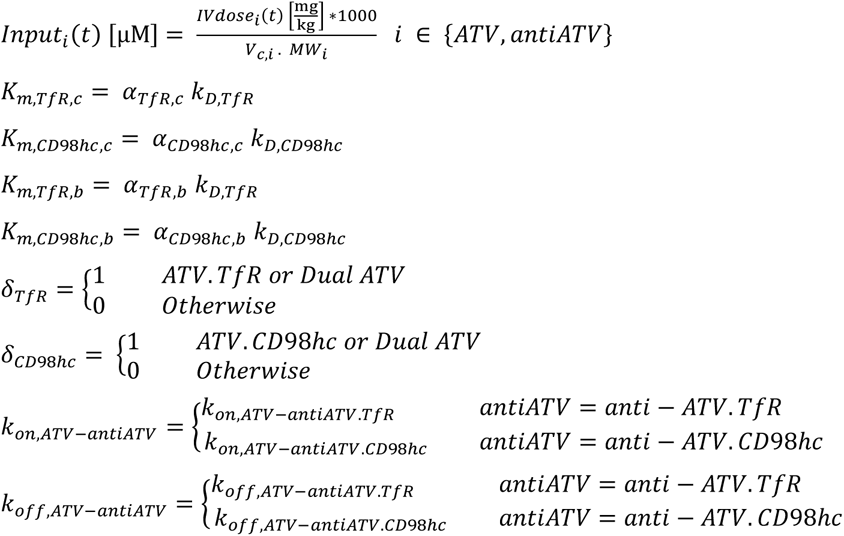

### Model-predicted observables corresponding to brain fluxes

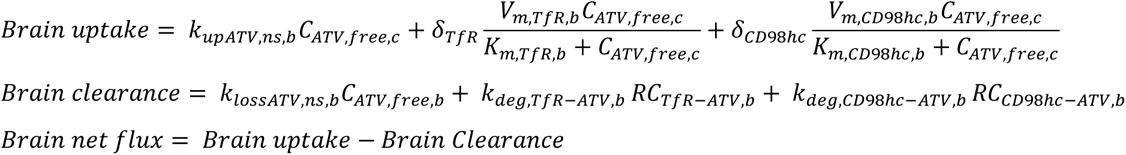

### Model-predicted observables corresponding to bioanalytical measurements

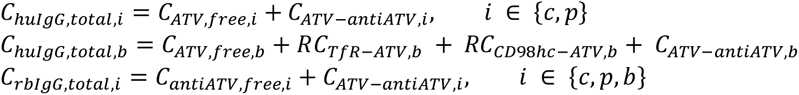

Where,

*C*ℎ*uIgG*,*total* and *CrbIgG*,*total* correspond to the model-predicted total human IgG (huIgG) and total rabbit IgG (rbIgG) assay measurements, respectively.

## Data Availability

Coordinates and related data for the crystal structures of the ATV.TfR**–**anti-ATV.TfR and ATV.CD98hc**–**anti-ATV.CD98hc complexes have been deposited in the PDB. Coordinates will be released upon final publication. PK model files will also be released upon final publication.

## Author Contributions

A.M., D.C., A.Q., M.S.K., K.G., R.G.T., C.M.K., and K.R.H. designed research; A.M., D.C., A.Q., J.E.K., K.S.C., R.C.W., J.C., R.T., N.P.D.L., J.D., P.A., A.B., I.B., B.B., E.Y.T.C., R.C., A.C., D.H., D.J.K., N.K., K.J.L., E.L., H.M., H.N., M.E.P., K.R, Y.R.-C, M.S., T.T, R.K.A.V., and K.X. performed research; A.A., M.S.D., J.W.L., C.S.M., L.R., J.M.S., R.J.W., D.Z., Y.J.Y.Z., S.D., R.G.T., C.M.K., and K.R.H. supervised research; A.M., D.C., A.Q., J.E.K., C.M.K., and K.R.H. wrote the paper.

