## Supplementary Information for "Flux analysis resolves distinct brain uptake and clearance of TfR- and CD98hc-targeting large molecules"

**This file includes:**

Supplementary text and Tables S1 to S8
Figs. S1 to S8
SI References

**Table S1. ATV molecules and experimental datasets included for PK model calibration and validation.**

| Calibration Dataset | | | | |
|---|---|---|---|---|
| **Molecule name** | **Binding affinity ($K_D$)** | **Mouse model** | **PK Matrix** | **Dose range (mg/kg)** |
| anti-DNP | – | $TfR^{mu/hu}$ KI<br>$CD98hc^{mu/hu}$ KI<br>$TfR^{mu/hu}$/$CD98hc^{mu/hu}$ DKI | Plasma<br>Brain | 10,15,50 |
| ATV.TfR100 | 100 nM | $TfR^{mu/hu}$ KI<br>$TfR^{mu/hu}$/$CD98hc^{mu/hu}$ DKI | Plasma<br>Brain | 30 |
| ATV.TfR110 | 110 nM | $TfR^{mu/hu}$ KI | Plasma<br>Brain | 10,50 |
| ATV.TfR620 | 620 nM | $TfR^{mu/hu}$ KI | Plasma<br>Brain | 10,25,<br>30,50 |
| ATV.TfR1100 | 1100 nM | $TfR^{mu/hu}$ KI | Plasma<br>Brain | 10, 50 |
| ATV.TfR2500 | 2500 nM | $TfR^{mu/hu}$ KI | Plasma<br>Brain | 10, 30 |
| ATV.TfR8000 | 8000 nM | $TfR^{mu/hu}$ KI | Plasma<br>Brain | 50 |
| ATV.CD98hc20 | 20 nM | $CD98hc^{mu/hu}$ KI | Plasma<br>Brain | 15 |
| ATV.CD98hc170 | 170 nM | $CD98hc^{mu/hu}$ KI<br>$TfR^{mu/hu}$/$CD98hc^{mu/hu}$ DKI | Plasma<br>Brain | 1,3,10,<br>15, 25,30,50 |
| ATV.CD98hc230 | 230 nM | $CD98hc^{mu/hu}$ KI | Plasma<br>Brain | 50 |
| ATV.CD98hc550 | 550 nM | $CD98hc^{mu/hu}$ KI<br>$TfR^{mu/hu}$/$CD98hc^{mu/hu}$ DKI | Plasma<br>Brain | 15,30 |
| ATV.CD98hc610 | 610 nM | $CD98hc^{mu/hu}$ KI | Plasma<br>Brain | 50 |
| ATV.CD98hc810 | 810 nM | $CD98hc^{mu/hu}$ KI | Plasma<br>Brain | 50 |
| ATV.CD98hc1400 | 1400 nM | $CD98hc^{mu/hu}$ KI | Plasma<br>Brain | 50 |
| ATV.CD98hc6300 | 6300 nM | $CD98hc^{mu/hu}$ KI<br>$TfR^{mu/hu}$/$CD98hc^{mu/hu}$ DKI | Plasma<br>Brain | 30,50 |
| ATV.TfR620/CD98hc170 | Table S2 | $TfR^{mu/hu}$/$CD98hc^{mu/hu}$ DKI | Plasma<br>Brain | 10,25,50 |

| Validation Dataset | | | | |
|---|---|---|---|---|
| **Molecule name** | **Binding affinity ($K_D$)** | **Mouse model** | **PK Matrix** | **Dose range (mg/kg)** |
| **ATV.TfR100/CD98hc170** | **Table S2** | **$TfR^{mu/hu}/CD98hc^{mu/hu}$ DKI** | **Plasma Brain** | **30** |
| **ATV.TfR100/CD98hc550** | **Table S2** | **$TfR^{mu/hu}/CD98hc^{mu/hu}$ DKI** | **Plasma Brain** | **30** |
| **ATV.TfR100/CD98hc6300** | **Table S2** | **$TfR^{mu/hu}/CD98hc^{mu/hu}$ DKI** | **Plasma Brain** | **30** |
| **ATV.TfR620/CD98hc170** | **Table S2** | **$TfR^{mu/hu}/CD98hc^{mu/hu}$ DKI** | **Plasma Brain** | **30** |
| **ATV.TfR620/CD98hc550** | **Table S2** | **$TfR^{mu/hu}/CD98hc^{mu/hu}$ DKI** | **Plasma Brain** | **30** |
| **ATV.TfR620/CD98hc6300** | **Table S2** | **$TfR^{mu/hu}/CD98hc^{mu/hu}$ DKI** | **Plasma Brain** | **30** |
| **ATV.TfR2500/CD98hc170** | **Table S2** | **$TfR^{mu/hu}/CD98hc^{mu/hu}$ DKI** | **Plasma Brain** | **30** |
| **ATV.TfR2500/CD98hc550** | **Table S2** | **$TfR^{mu/hu}/CD98hc^{mu/hu}$ DKI** | **Plasma Brain** | **30** |
| **ATV.TfR2500/CD98hc6300** | **Table S2** | **$TfR^{mu/hu}/CD98hc^{mu/hu}$ DKI** | **Plasma Brain** | **30** |

**Table S2. Summary of binding affinity and kinetic parameters of dual ATV molecules.**

| Molecule name | CD98hc | | | | TfR | | | |
|---|---|---|---|---|---|---|---|---|
| | $K_D$ (nM) | | $k_{on}$ ($M^{-1}s^{-1}$) | $k_{off}$ ($s^{-1}$) | $K_D$ (nM) | | $k_{on}$ ($M^{-1}s^{-1}$) | $k_{off}$ ($s^{-1}$) |
| | Expected | Actual | | | Expected | Actual | | |
| **ATV.TfR100/CD98hc170** | 170 | 150 | 4.05e4 | 5.97e-3 | 100 | 200 | 2.79e5 | 5.61e-2 |
| **ATV.TfR100/CD98hc550** | 550 | 820 | 5.54e4 | 4.57e-2 | 100 | 200 | 2.75e5 | 5.39e-2 |
| **ATV.TfR100/CD98hc6300** | 6300 | 7400 | 5.70e4 | 4.23e-1 | 100 | 200 | 2.76e5 | 5.64e-2 |
| **ATV.TfR620/CD98hc170** | 170 | 150 | 3.94e4 | 5.89e-3 | 620 | 830 | 1.88e5 | 1.57e-1 |
| **ATV.TfR620/CD98hc550** | 550 | 680 | 5.86e4 | 3.96e-2 | 620 | 750 | 1.54e5 | 1.16e-1 |
| **ATV.TfR620/CD98hc6300** | 6300 | 6600 | 5.41e4 | 3.55e-1 | 620 | 810 | 1.89e5 | 1.53e-1 |
| **ATV.TfR2500/CD98hc170** | 170 | 130 | 4.38e4 | 5.84e-3 | 2500 | 5100 | 4.81e4 | 2.45e-1 |
| **ATV.TfR2500/CD98hc550** | 550 | 780 | 5.74e4 | 4.47e-2 | 2500 | 4900 | 5.19e4 | 2.54e-1 |
| **ATV.TfR2500/CD98hc6300** | 6300 | 5700 | 5.38e4 | 3.07e-1 | 2500 | 6000 | 4.77e4 | 2.86e-1 |

Expected affinities are derived from the average affinity of the corresponding monovalent TV molecules, whereas actual affinities represent binding of the dual ATV molecules to the TfR apical domain and CD98hc extracellular domain (ECD) (1).

**Table S3. ATV PK model state variables.**

| State Variable | Model Module | Description | Units |
|---|---|---|---|
| $\boldsymbol{C_{ATV,free,i}}$ | ATV PK model | Concentration of free ATV in compartment $i$,<br>where $i \in \{c,p,b\}$ denotes the central, peripheral, and brain compartments.<br>ATV represents ATV.TfR, ATV.CD98hc, or dual ATV. | μM |
| $\boldsymbol{R_{TfR,free,b}}$ | ATV PK model | Concentration of free transferrin receptor (TfR) in the brain compartment. | μM |
| $\boldsymbol{R_{CD98hc,free,b}}$ | ATV PK model | Concentration of free CD98hc receptor in the brain compartment. | μM |
| $\boldsymbol{RC_{TfR-ATV,b}}$ | ATV PK model | Concentration of the TfR–ATV receptor complex in the brain compartment.<br>ATV represents either ATV.TfR or dual ATV. | μM |
| $\boldsymbol{RC_{CD98hc-ATV,b}}$ | ATV PK model | Concentration of the CD98hc–ATV receptor complex in the brain compartment.<br>ATV represents either ATV.CD98hc or dual ATV. | μM |
| $\boldsymbol{C_{antiATV,free,i}}$ | ATV–antiATV Interaction | Concentration of free anti-ATV antibody in compartment $i$,<br>where $i \in \{c,p,b\}$ denotes the central, peripheral, and brain compartments.<br>antiATV represents either anti-ATV.TfR or anti-ATV.CD98hc. | μM |
| $\boldsymbol{C_{ATV-antiATV,i}}$ | ATV–antiATV Interaction | Concentration of bound ATV–antiATV complex in compartment I,<br>where $i \in \{c,p,b\}$ denotes the central, peripheral, and brain compartments.<br>ATV–antiATV represents either ATV.TfR–anti-ATV.TfR<br>or ATV.CD98hc–anti-ATV.CD98hc. | μM |

**Table S4. ATV PK model parameters.**

| Parameter | Description | Value (CV%) | Units | Source |
|---|---|---|---|---|
| $V_{c,ATV}$ | Central volume of distribution of ATV | 42.03 (0.27) | mL/kg | Model fit |
| $V_{p,ATV}$ | Peripheral volume of distribution of ATV | 84.31 (3.42) | mL/kg | Model fit |
| $CL_{ATV}$ | Non-specific systemic clearance of ATV | 0.098 (8.06) | mL/hr/kg | Model fit |
| $CL_{D,ATV}$ | Intercompartmental distribution clearance of ATV | 2.47 (7.13) | mL/hr/kg | Model fit |
| $V_{m,TfR,c}$ | Maximal TfR-mediated systemic clearance | 0.043 (3.89) | µM/hr | Model fit |
| $K_{D,TfR}$ | Binding affinity of ATV to TfR | Molecule specific | µM | Table S1 |
| $\alpha_{TfR,c}$ | Michaelis-Menten constant scaling factor for $K_{m,TfR,c}$ | 0.76 (10.44) | Dimensionless | Model fit |
| $K_{m,TfR,c}$ | Michaelis-Menten constant for systemic TfR-mediated clearance | $\alpha_{TfR,c}\ K_{D,TfR}$ | µM | - |
| $V_{m,CD98hc,c}$ | Maximal systemic CD98hc-mediated clearance | 0.027 (6.97) | µM/hr | Model fit |
| $K_{D,CD98hc}$ | Binding affinity of ATV to CD98hc | Molecule specific | µM | Table S1 |
| $\alpha_{CD98hc,c}$ | Scaling factor for $K_{m,CD98hc,c}$ | 5.18 (9.44) | Dimensionless | Model fit |
| $K_{m,CD98hc,c}$ | Michaelis-Menten constant for systemic CD98hc-mediated clearance | $\alpha_{CD98hc,c} \cdot K_{D,CD98hc}$ | µM | - |
| $k_{up,ATV,ns,b}$ | Non-specific ATV brain uptake rate constant | 6.39e-5 (15.15) | 1/hr | Model fit |
| $k_{lossATV,ns,b}$ | Non-specific ATV brain loss rate constant | 0.029 (17.22) | 1/hr | Model fit |
| $V_{m,TfR,b}$ | Maximal TfR-mediated brain uptake | 0.0034 (6.96) | µM/hr | Model fit |
| $\alpha_{TfR,b}$ | Michaelis-Menten constant scaling factor for $K_{m,TfR,b}$ | 3.25 (13.06) | Dimensionless | Model fit |
| $R_{TfR,b}$ | Brain TfR concentration | 0.59 (21.38) | µM | Model fit |
| $V_{m,CD98hc,b}$ | Maximal CD98hc-mediated brain uptake | 0.00019 (7.74) | µM/hr | Model fit |
| $\alpha_{CD98hc,b}$ | Michaelis-Menten constant scaling factor for $K_{m,CD98hc,b}$ | Same as $\alpha_{CD98hc,c}$ | Dimensionless | - |
| $R_{CD98hc,b}$ | Brain CD98hc receptor concentration | 3.34 (13.41) | µM | Model fit |
| $k_{deg,TfR}$ | TfR degradation rate | 8.40 | 1/hr | Pardridge et al. 2021(2) |
| $k_{syn,TfR}$ | TfR synthesis rate | $k_{deg,TfR} \cdot R_{TfR,b}$ | µM/hr | - |

| | | | | |
|---|---|---|---|---|
| $\mathbf{k_{on,TfR}}$ | Association rate constant for ATV binding to TfR | 600 | 1/µM/hr | Average from Table S2 |
| $\mathbf{k_{off,TfR}}$ | Dissociation rate constant for TfR-bound ATV complex | $K_{D,TfR} \cdot k_{on,TfR}$ | 1/hr | - |
| $\mathbf{k_{deg,TfR–ATV,b}}$ | Degradation rate of ATV bound to TfR in brain | 0.14 (8.21) | 1/hr | Model fit |
| $\mathbf{k_{deg,CD98hc}}$ | CD98hc degradation rate | 0.0044 (10.49) | 1/hr | Model fit |
| $\mathbf{k_{syn,CD98hc}}$ | CD98hc synthesis rate | $k_{deg,CD98hc} \cdot R_{CD98hc,b}$ | µM/hr | - |
| $\mathbf{k_{on,CD98hc}}$ | Association rate constant for ATV binding to CD98hc | 184 | 1/µM/hr | Average from Table S2 |
| $\mathbf{k_{off,CD98hc}}$ | Dissociation rate constant for CD98hc-bound ATV complex | $K_{D,CD98hc} \cdot k_{on,CD98hc}$ | 1/hr | - |
| $\mathbf{k_{deg,CD98hc–ATV,b}}$ | Degradation rate of ATV bound to CD98hc in brain | Same as $\mathbf{k_{deg,CD98hc}}$ | 1/hr | Model fit |
| $\mathbf{MW_{ATV}}$ | Molecular weight of ATV | 146 | kDa | - |

**Table S5. PK model parameters for anti-ATV antibodies and ATV–anti-ATV interactions.**

| Parameter | Description | Value (CV%) | Units | Source |
|---|---|---|---|---|
| $\mathbf{V_{c,antiATV}}$ | Central volume of distribution of antiATV | same as $V_{c,ATV}$ | mL/kg | Table S4 |
| $\mathbf{V_{p,antiATV}}$ | Peripheral volume of distribution of antiATV | same as $V_{p,ATV}$ | mL/kg | Table S4 |
| $\mathbf{CL_{antiATV}}$ | Non-specific systemic clearance of antiATV | same as $CL_{ATV}$ | mL/hr/kg | Table S4 |
| $\mathbf{CL_{D,antiATV}}$ | Intercompartmental distribution clearance of antiATV | same as $CL_{D,ATV}$ | mL/hr/kg | Table S4 |
| $\mathbf{k_{on,ATV\text{-}antiATVTfR}}$ | Association rate constant for antiATV.TfR binding to ATV.TfR | 14400 | 1/µM/hr | Average from Table S6 |
| $\mathbf{k_{D,ATV–antiATVTfR}}$ | Binding affinity of antiATV.TfR to ATV.TfR | 2.80e-7 | µM | Average from Table S6 |
| $\mathbf{k_{off,ATV–antiATVTfR}}$ | Dissociation rate constant for antiATV.TfR-ATV.TfR complex | $K_{D,ATV–antiATVTfR} \cdot k_{on,ATV–antiATVTfR}$ | 1/hr | – |
| $\mathbf{k_{on,ATV–antiATVCD98hc}}$ | Association rate constant for antiATV.CD98hc binding to ATV.CD98hc | 85 | 1/µM/hr | Average from Table S6 |
| $\mathbf{k_{D,ATV–antiATVCD98hc}}$ | Binding affinity of antiATV.CD98hc to ATV.CD98hc | 0.00174 | µM | Average from Table S6 |
| $\mathbf{k_{off,ATV–antiATVCD98hc}}$ | Dissociation rate constant for antiATV.CD98hc-ATV.CD98hc complex | $K_{D,ATV–antiATVCD98hc} \cdot k_{on,ATV–antiATVCD98hc}$ | 1/hr | – |
| $\mathbf{k_{up,antiATV,ns,b}}$ | Non-specific antiATV brain uptake rate constant, with antiATV representing either anti-ATV.TfR or anti-ATV.CD98hc. | same as $k_{up,ATV,ns,b}$ | 1/hr | Table S4 |
| $\mathbf{k_{loss,antiATV,ns,b}}$ | Non-specific antiATV brain loss rate constant, with antiATV representing either anti-ATV.TfR or anti-ATV.CD98hc. | same as $k_{loss,ATV,ns,b}$ | 1/hr | Table S4 |
| $\mathbf{k_{deg,ATV–antiATV,b}}$ | Degradation rate of ATV-antiATV complex in brain | same as $k_{loss,ATV,ns,b}$ | 1/hr | Table S4 |
| $\mathbf{MW_{antiATV}}$ | antiATV molecular weight | 146 | kDa | Table S4 |

**Table S6. Binding affinities and kinetic parameters of anti-ATV antibodies for ATV molecules determined by KinExA.**

| anti-ATV.CD98hc | ATV.CD98hc170 | ATV.CD98hc550 | ATV.CD98hc6300 |
|---|---|---|---|
| $K_D$ (nM) | 0.5<br>(0.2-1.1) | 1.22<br>(0.7-1.9) | 3.5<br>(2.4-5.1) |
| $k_{off}$ ($s^{-1}$) | $1.4 \times 10^{-5}$ | $2.8 \times 10^{-5}$ | $6.8 \times 10^{-5}$ |
| $k_{on}$ ($M^{-1}s^{-1}$) | $2.8 \times 10^{4}$<br>($2.5 \times 10^{4} – 3.1 \times 10^{4}$) | $2.3 \times 10^{4}$<br>($1.6 \times 10^{4} – 3.0 \times 10^{4}$) | $1.94 \times 10^{4}$<br>($1.80 \times 10^{4} – 2.1 \times 10^{4}$) |

| anti-ATV.TfR | ATV.TfR100 | ATV.TfR620 | ATV.TfR2500 |
|---|---|---|---|
| $K_D$ (fM) | 100<br>(56.3-181) | 515<br>(291-793) | 226<br>(68-494) |
| $k_{off}$ ($s^{-1}$) | $4.9 \times 10^{-7}$ | $1.7 \times 10^{-6}$ | $9.7 \times 10^{-7}$ |
| $k_{on}$ ($M^{-1}s^{-1}$) | $4.4 \times 10^{6}$<br>($3.4 \times 10^{6} – 5.9 \times 10^{6}$) | $3.3 \times 10^{6}$<br>($2.8 \times 10^{6} – 3.8 \times 10^{6}$) | $4.3 \times 10^{6}$<br>($3.8 \times 10^{6} – 4.8 \times 10^{6}$) |

Bracketed numbers represent 95% confidence interval of the measurements.

**Estimation of apparent brain loss rate constants ($k_{loss}$) corresponding to Fig 4G.** The post–anti-ATV phase of the brain PK profiles of Figure 4C,D was fitted to estimate the apparent first-order brain loss rate constant ($k_{loss}$). For ATV.CD98hc variants (Figure 4D), the post–anti-ATV.CD98hc brain concentrations were described by a mono-exponential decay model,

$$C(t) = C_0 e^{-k_{\mathrm{loss}} t},$$

where $C_0$ denotes the concentration at the onset of the decay phase.

For ATV.TfR variants (Figure 4C), the post–anti-ATV.TfR brain PK profiles exhibited a low-level terminal plateau. These profiles were therefore fitted using a mono-exponential decay model with a residual concentration term,

$$C(t) = C_0 e^{-k_{\mathrm{loss}} t} + C_{\mathrm{res}},$$

where $C_{res}$ represents the residual concentration during the terminal phase of the post–anti-ATV brain PK profile. Accordingly, $k_{loss}$ describes the apparent first-order loss rate constant of the decaying component above the residual plateau.

Fits to the observed data were performed by nonlinear least-squares regression using the *lsqcurvefit* function, and parameter uncertainty was quantified by calculating 90% confidence intervals from the estimated Jacobian matrix using the *nlparci* function (MATLAB 2024b; MathWorks). Fit results are provided in Table S7.

The corresponding model-derived $k_{loss}$ estimates (solid and dashed traces in Figure 4G) were estimated from the simulated brain PK profiles following anti-ATV administration using the same fitting functions and estimation procedure applied to the experimental data, thereby ensuring a consistent comparison between model predictions and pulse–chase measurements.

**Table S7 – Mono-exponential fit parameters used to estimate apparent brain loss rate constants ($k_{loss}$) from the pulse-chase experiments.**

| Experiment | $C_0$ (nM) [90% CI] | $k_{loss}$ (1/h) [90% CI] | $C_{res}$ (nM) [90% CI] |
|---|---|---|---|
| **anti-ATV.TfR + ATV.TfR100** | 17.17 [16.03, 18.31] | 0.073 [0.059, 0.088] | 2.04 [1.36, 2.72] |
| **anti-ATV.TfR + ATV.TfR620** | 11.23 [10.01, 12.45] | 0.062 [0.042, 0.082] | 1.86 [1.09, 2.64] |
| **anti-ATV.TfR + ATV.TfR2500** | 4.34 [3.78, 4.91] | 0.055 [0.033, 0.076] | 1.34 [0.96, 1.73] |
| **anti-ATV.CD98hc + ATV.CD98hc170** | 19.34 [17.73, 20.94] | 0.0042 [0.0032, 0.0051] | – |
| **anti-ATV.CD98hc + ATV.CD98hc550** | 11.99 [10.32, 13.66] | 0.0051 [0.0031,0.0072] | – |
| **anti-ATV.CD98hc + ATV.CD98hc6300** | 4.18 [3.41, 4.95] | 0.0050 [0, 0.01] | – |

**Table S8 – X-ray crystallography data collection and refinement statistics for anti-ATV Fab complexes.**

| **PDB ID** | **ATV.TfR–anti-ATV.TfR**<br>PDB ID and coordinates will be released upon journal publication | **ATV.CD98hc–anti-ATV.CD98hc**<br>PDB ID and coordinates will be released upon journal publication |
|---|---|---|
| **Data collection** | | |
| Space group | P 21 21 21 | P 41 21 2 |
| Cell dimensions | | |
| *a*, *b*, *c* (Å) | 62.1, 79.5, 217.4 | 99.7, 99.7, 205.1 |
| a, b, g (°) | 90.0, 90.0, 90.0 | 90.0, 90.0, 90.0 |
| Resolution (Å) | 108.94-2.00 (2.04-2.00) | 45.64-2.71 (2.84-2.71) |
| $R_{sym}$ or $R_{merge}$ | 0.142 (3.579) | 0.147 (1.167) |
| *I* / s*I* | 9.8 (0.9) | 15.2 (2.5) |
| $CC_{1/2}$ (%) | 0.998 (0.412) | 0.998 (0.774) |
| Completeness (%) | 99.9 (97.7) | 100.0 (100.0) |
| Redundancy | 13.4 (13.8) | 12.7 (12.6) |
| **Refinement** | | |
| Resolution (Å) | 2.00 | 2.71 |
| No. unique reflections | 73798 | 28914 |
| $R_{work}$ / $R_{free}$ | 0.229/0.273 | 0.232/0.276 |
| No. atoms | | |
| Protein | 6542 | 6445 |
| Ligand/ion | 133 | 172 |
| Water | 218 | 17 |
| *B*-factors | | |
| Protein | 64.48 | 70.43 |
| Ligand/ion | 107.27 | 135.67 |
| Water | 50.46 | 40.3 |
| R.m.s. deviations | | |
| Bond lengths (Å) | 0.0143 | 0.0074 |
| Bond angles (°) | 2.549 | 1.758 |

Values in parentheses correspond to the highest-resolution shell.

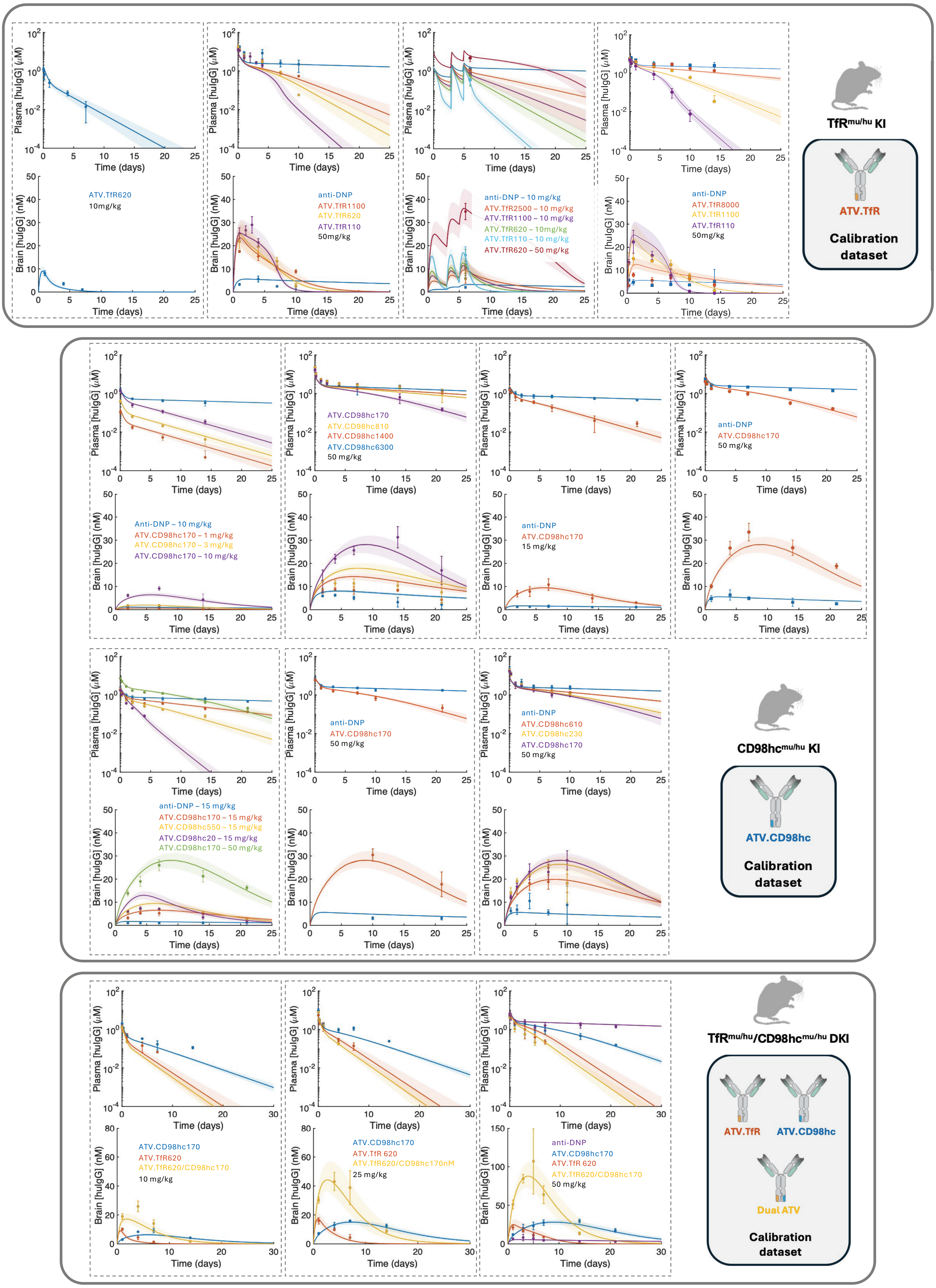

Plasma [huIgG] (μM)
Brain [huIgG] (nM)
Time (days)
ATV.TfR620
10mg/kg
anti-DNP
ATV.TfR1100
ATV.TfR620
ATV.TfR110
50mg/kg
anti-DNP – 10 mg/kg
ATV.TfR2500 – 10 mg/kg
ATV.TfR1100 – 10 mg/kg
ATV.TfR620 – 10mg/kg
ATV.TfR110 – 10 mg/kg
ATV.TfR620 – 50 mg/kg
anti-DNP
ATV.TfR8000
ATV.TfR1100
ATV.TfR110
50mg/kg
TfRmu/hu KI
ATV.TfR
Calibration dataset
Anti-DNP – 10 mg/kg
ATV.CD98hc170 – 1 mg/kg
ATV.CD98hc170 – 3 mg/kg
ATV.CD98hc170 – 10 mg/kg
ATV.CD98hc170
ATV.CD98hc810
ATV.CD98hc1400
ATV.CD98hc6300
50 mg/kg
anti-DNP
ATV.CD98hc170
15 mg/kg
anti-DNP
ATV.CD98hc170
50 mg/kg
anti-DNP – 15 mg/kg
ATV.CD98hc170 – 15 mg/kg
ATV.CD98hc550 – 15 mg/kg
ATV.CD98hc20 – 15 mg/kg
ATV.CD98hc170 – 50 mg/kg
anti-DNP
ATV.CD98hc170
50 mg/kg
anti-DNP
ATV.CD98hc610
ATV.CD98hc230
ATV.CD98hc170
50 mg/kg
CD98hcmu/hu KI
ATV.CD98hc
Calibration dataset
ATV.CD98hc170
ATV.TfR620
ATV.TfR620/CD98hc170
10 mg/kg
ATV.CD98hc170
ATV.TfR 620
ATV.TfR620/CD98hc170nM
25 mg/kg
anti-DNP
ATV.CD98hc170
ATV.TfR 620
ATV.TfR620/CD98hc170
50 mg/kg
TfRmu/hu/CD98hcmu/hu DKI
ATV.TfR
ATV.CD98hc
Dual ATV
Calibration dataset

**Fig. S1. Visual predictive checks of the ATV PK model using the calibration dataset.** Plasma and brain PK profiles comprising the calibration datasets used for development of the PK models for ATV.TfR (top), ATV.CD98hc (middle), and dual ATV (bottom). Calibration datasets span a broad range of receptor binding affinities and dose levels. ATV.TfR studies were performed in $TfR^{mu/hu}$ KI mice, ATV.CD98hc studies in $CD98hc^{mu/hu}$ KI mice, and dual ATV studies in $TfR^{mu/hu}/CD98hc^{mu/hu}$ DKI mice. Symbols represent the observed data (presented as mean ± SD), solid lines represent model fits, and shaded regions denote the 90% prediction intervals based on the uncertainty in the estimated model parameters (%CV in Table S4). Simulations of the dual ATV clones are done with the actual affinity values reported in Table S2. Note: Raw data for ATV.TfR620, ATV.CD98hc170, and ATV.TfR620/CD98hc170 at 25 mg/kg in $TfR^{mu/hu}/CD98hc^{mu/hu}$ DKI mice were previously published by Wells et al (1). Some of the PK data for ATV.CD98hc molecules were also previously published by Chew et al (3).

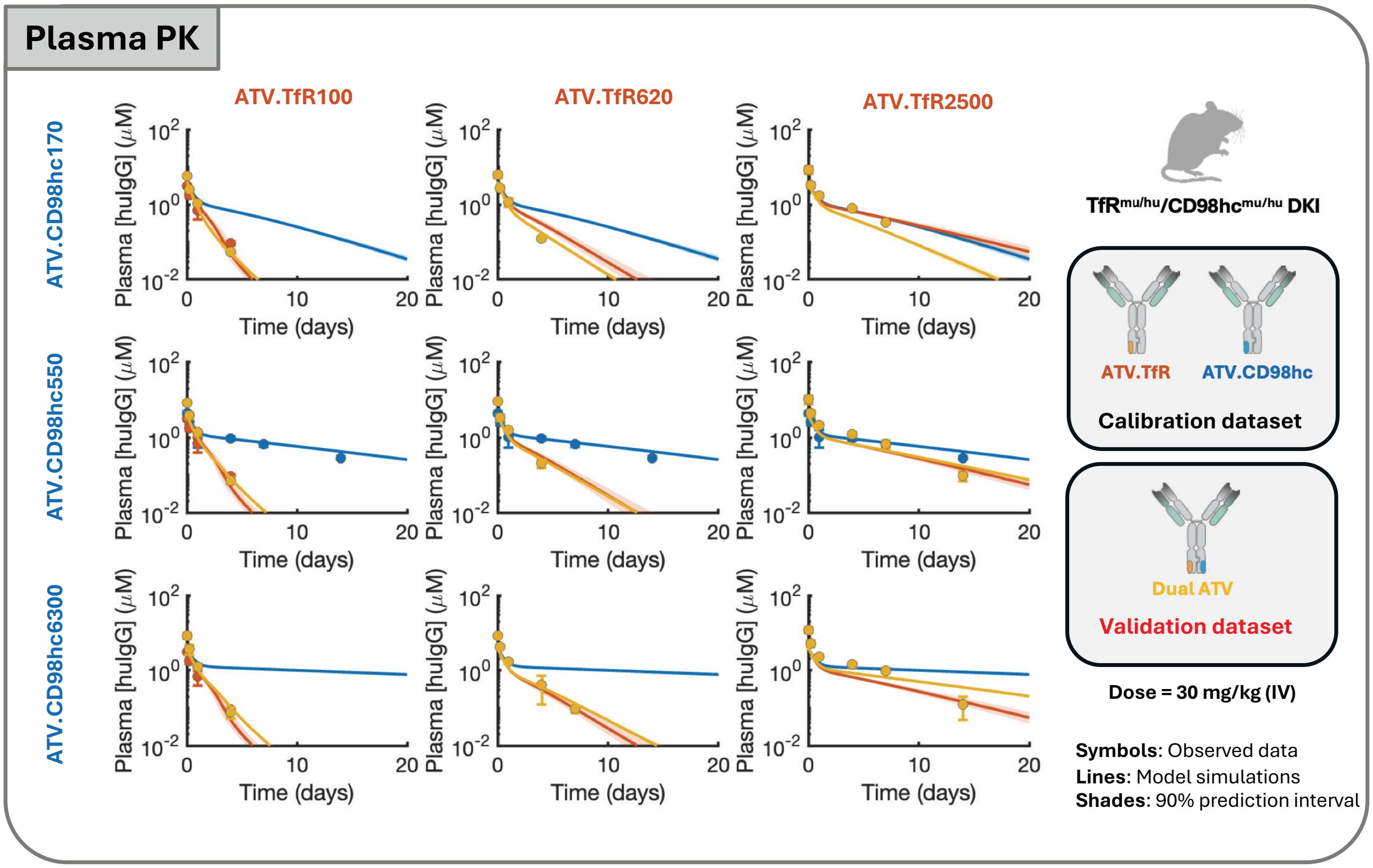

Plasma PK
ATV.TfR100
ATV.TfR620
ATV.TfR2500
ATV.CD98hc170
ATV.CD98hc550
ATV.CD98hc6300
Plasma [huIgG] ($\mu$M)
$10^{2}$
$10^{0}$
$10^{-2}$
0
10
20
Time (days)
TfR$^{mu/hu}$/CD98hc$^{mu/hu}$ DKI
ATV.TfR
ATV.CD98hc
Calibration dataset
Dual ATV
Validation dataset
Dose = 30 mg/kg (IV)
**Symbols**: Observed data
**Lines**: Model simulations
**Shades**: 90% prediction interval


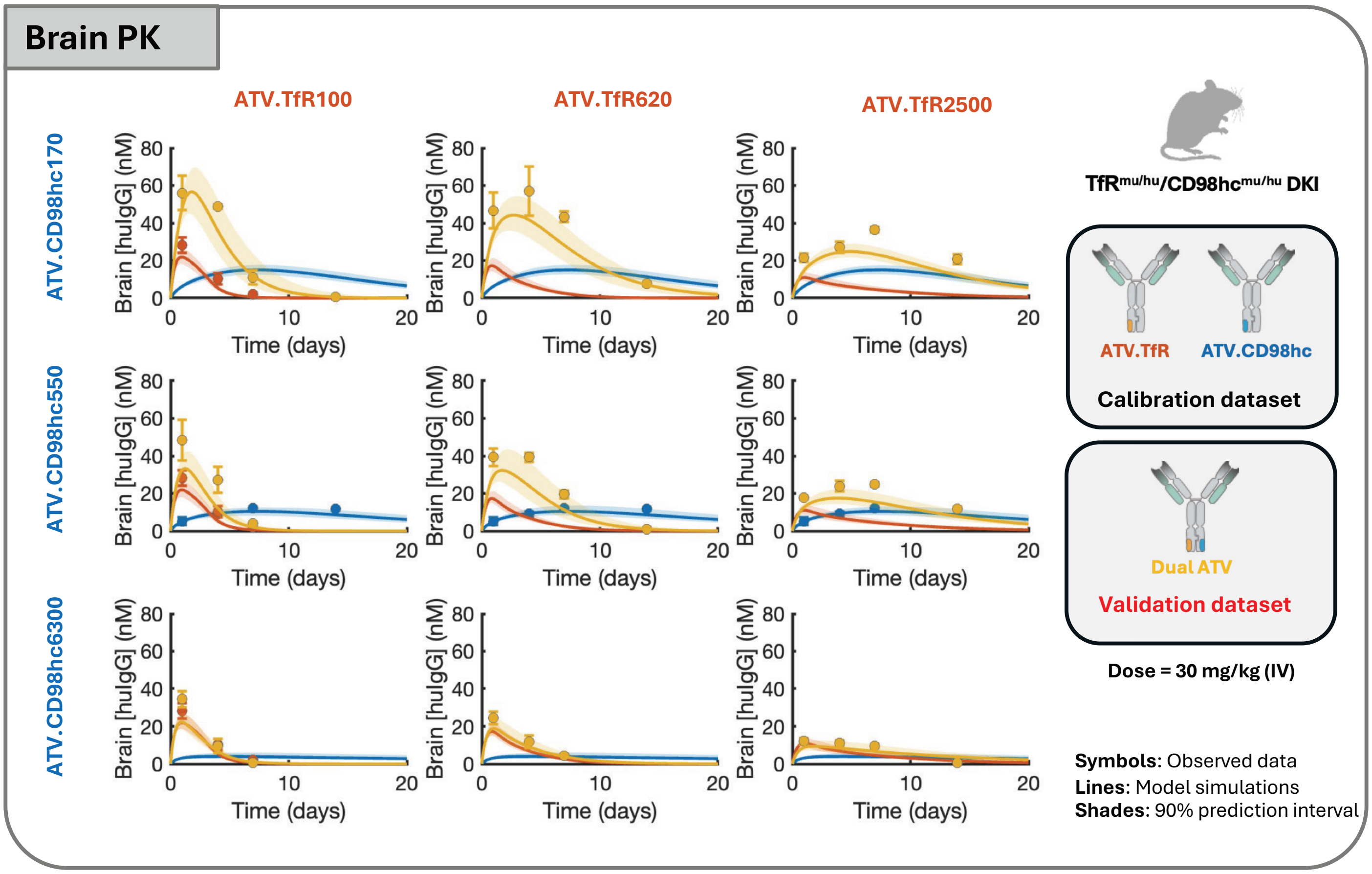

Brain PK
ATV.TfR100
ATV.TfR620
ATV.TfR2500
ATV.CD98hc170
ATV.CD98hc550
ATV.CD98hc6300
Brain [huIgG] (nM)
0
20
40
60
80
0
10
20
Time (days)
TfR$^{mu/hu}$/CD98hc$^{mu/hu}$ DKI
ATV.TfR
ATV.CD98hc
Calibration dataset
Dual ATV
Validation dataset
Dose = 30 mg/kg (IV)
**Symbols**: Observed data
**Lines**: Model simulations
**Shades**: 90% prediction interval

**Fig. S2. Calibration and Validation of the ATV PK model using a dual ATV affinity matrix.** Plasma (top) and brain (bottom) PK profiles of nine dual ATV constructs spanning a 3 × 3 matrix of TfR ($K_D$ = 100, 620, 2500 nM) and CD98hc ($K_D$ = 170, 550, 6300 nM) binding affinities following a single 30 mg/kg intravenous dose in $TfR^{mu/hu}$/$CD98hc^{mu/hu}$ DKI mice. Single-receptor ATV.TfR and ATV.CD98hc datasets were used for model calibration, whereas the dual ATV dataset was reserved for model validation. Symbols represent the observed data (presented as mean ± SD), solid lines represent model predictions, and shaded regions denote the 90% prediction intervals based on the uncertainty in the estimated model parameters (%CV in Table S4). Simulations of the dual ATV clones are done with the actual affinity values reported in Table S2. Raw data have also been previously published by Wells et al. (1)

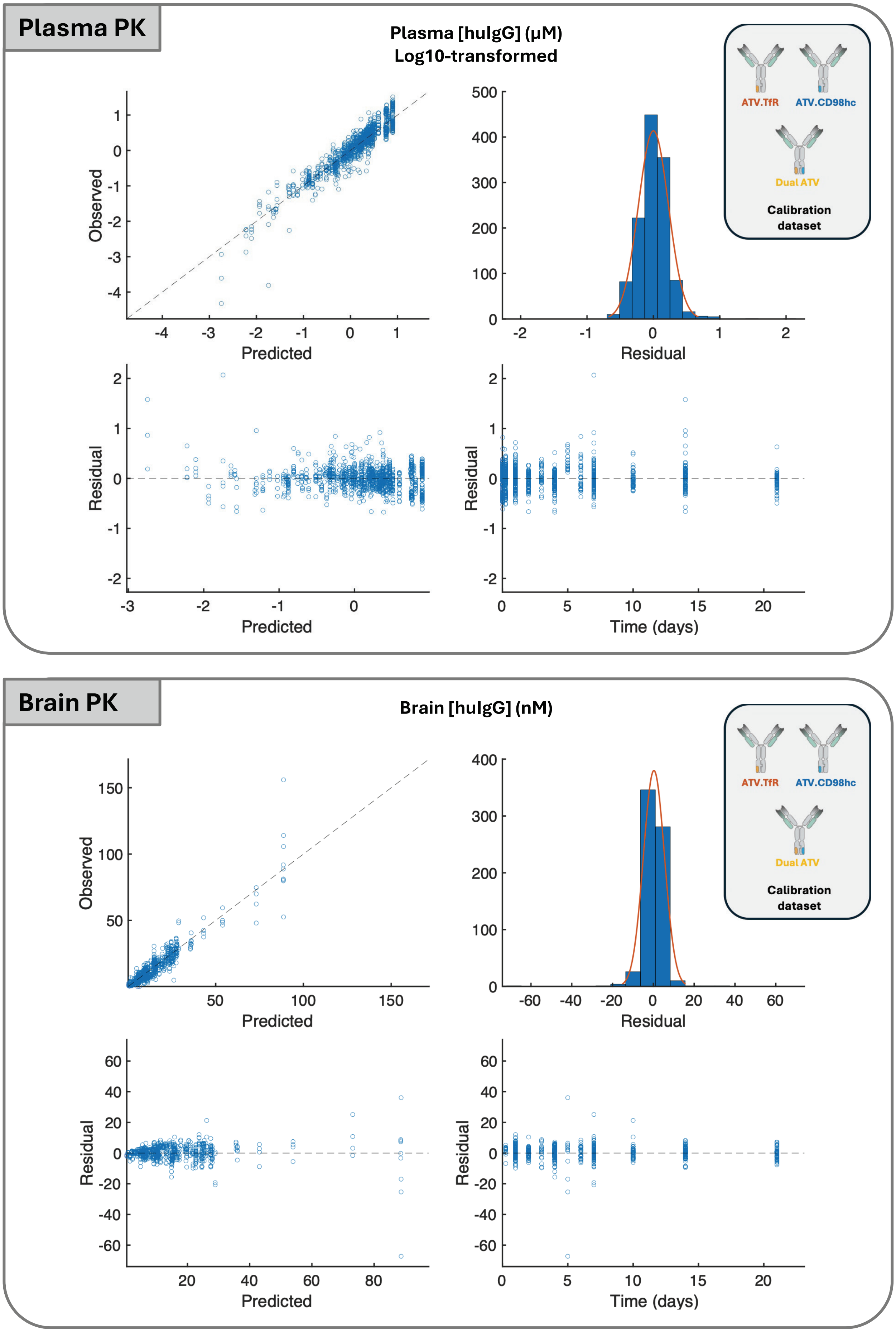
Plasma PK
Plasma [huIgG] (μM)
Log10-transformed
Observed
Predicted
Residual
Residual
Predicted
Residual
Time (days)
ATV.TfR
ATV.CD98hc
Dual ATV
Calibration dataset
Brain PK
Brain [huIgG] (nM)
Observed
Predicted
Residual
Residual
Predicted
Residual
Time (days)
ATV.TfR
ATV.CD98hc
Dual ATV
Calibration dataset

**Fig. S3. Goodness-of-fit diagnostics for the calibrated PK models using the calibration datasets.** Goodness-of-fit (GOF) diagnostics for plasma (top) and brain (bottom) PK data from the calibration datasets for ATV.TfR, ATV.CD98hc, and dual ATV. Panels show observed versus model-predicted concentrations, residual distributions, residuals versus model-predicted concentrations, and residuals versus time. Plasma concentrations were analyzed on the log10-transformed scale. Residuals are defined as observed minus model-predicted concentrations.

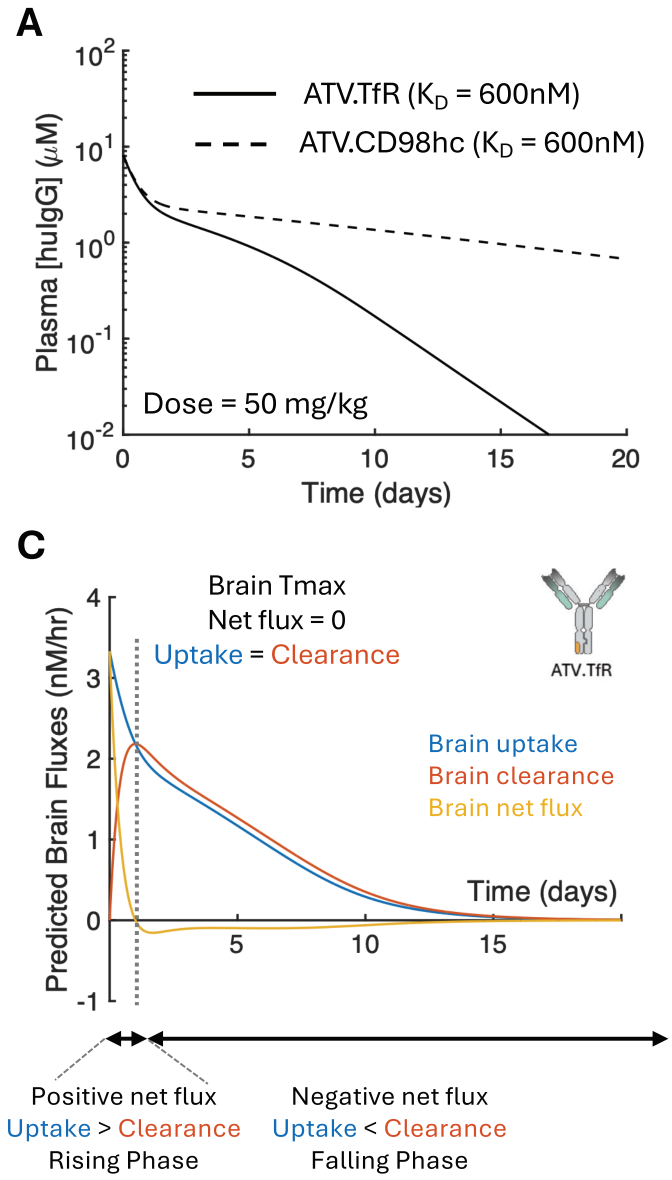
A
B
C
D
Plasma [huIgG] (μM)
Brain [huIgG] (nM)
Time (days)
ATV.TfR ($K_D$ = 600nM)
ATV.CD98hc ($K_D$ = 600nM)
Dose = 50 mg/kg
Predicted Brain Fluxes (nM/hr)
Brain Tmax
Net flux = 0
Uptake = Clearance
ATV.TfR
ATV.CD98hc
Brain uptake
Brain clearance
Brain net flux
Positive net flux
Uptake > Clearance
Rising Phase
Negative net flux
Uptake < Clearance
Falling Phase

**Fig. S4. Model-predicted brain fluxes of ATV.TfR and ATV.CD98hc at a matched receptor binding affinity. (A,B)** Simulated plasma (A) and brain (B) PK profiles of ATV.TfR and ATV.CD98hc with matched receptor binding affinity ($K_D$ = 600 nM) following a single 50 mg/kg intravenous dose. **(C,D)** Model-predicted brain uptake, brain clearance, and net brain fluxes for ATV.TfR (C) and ATV.CD98hc (D). Brain concentrations reach $T_{max}$ when brain uptake equals brain clearance (net brain flux = 0). Before $T_{max}$, brain uptake exceeds brain clearance, resulting in increasing brain concentrations, whereas after $T_{max}$, brain clearance exceeds brain uptake, resulting in declining brain concentrations.

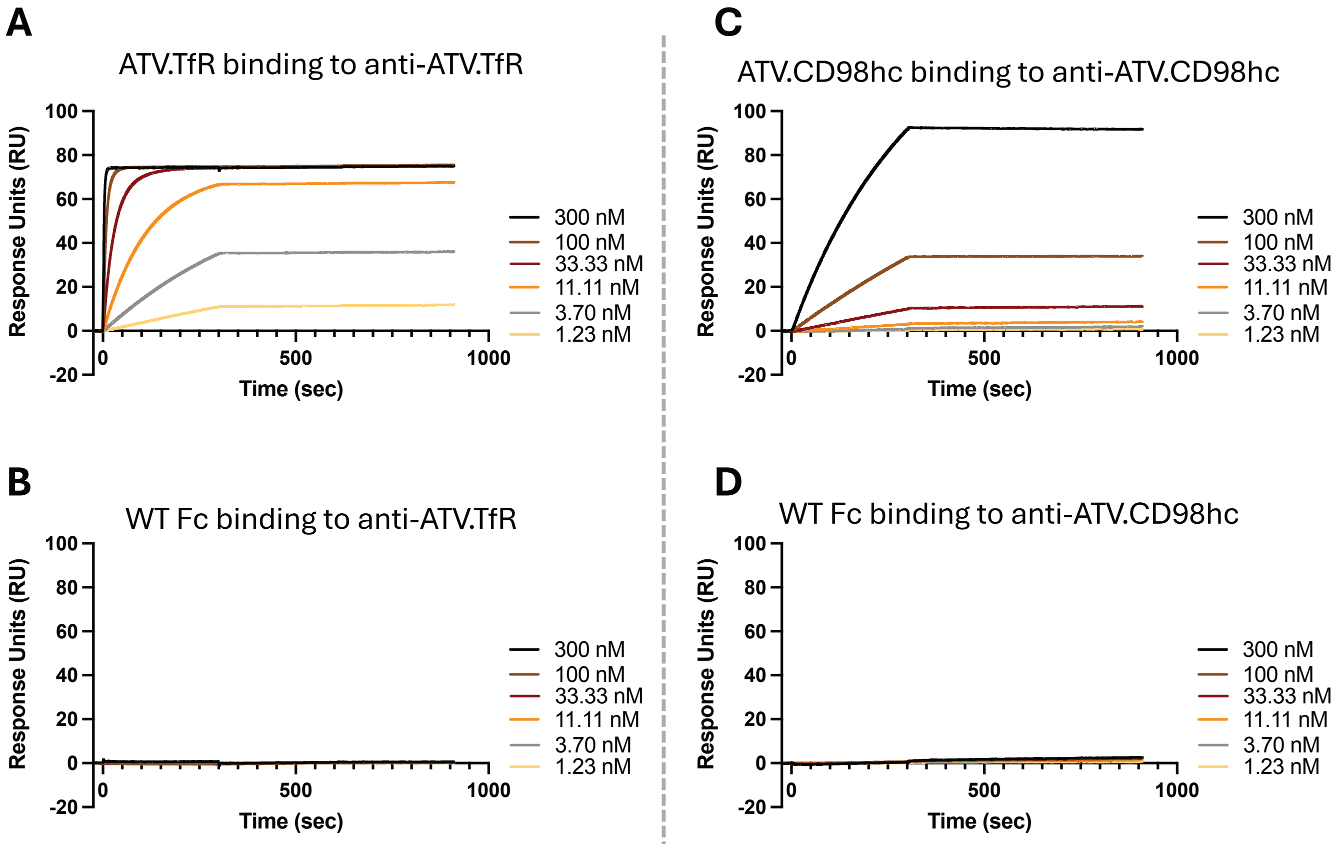
A
ATV.TfR binding to anti-ATV.TfR
B
WT Fc binding to anti-ATV.TfR
C
ATV.CD98hc binding to anti-ATV.CD98hc
D
WT Fc binding to anti-ATV.CD98hc
Response Units (RU)
Time (sec)
300 nM
100 nM
33.33 nM
11.11 nM
3.70 nM
1.23 nM

**Fig. S5. SPR measurements demonstrating no binding of anti-ATV.TfR and anti-ATV.CD98hc to the wild-type (WT) Fc. (A,B)** ATV.TfR (A) or human IgG1 (B) immobilized on anti-huFab CM5 sensor with Fab anti-ATV.TfR injected at a range of concentrations (300-1.23nM) for 300 sec and dissociated for 600 sec. **(C,D)** Fab anti-ATV.CD98hc was immobilized on anti-huFab CM5 sensor with Fc-only ATV.CD98hc (C) or human IgG1 (D) injected at a range of concentrations (300-1.23nM) for 300 sec and dissociated for 600 sec. Data was collected (n=2) using Biacore 8K.

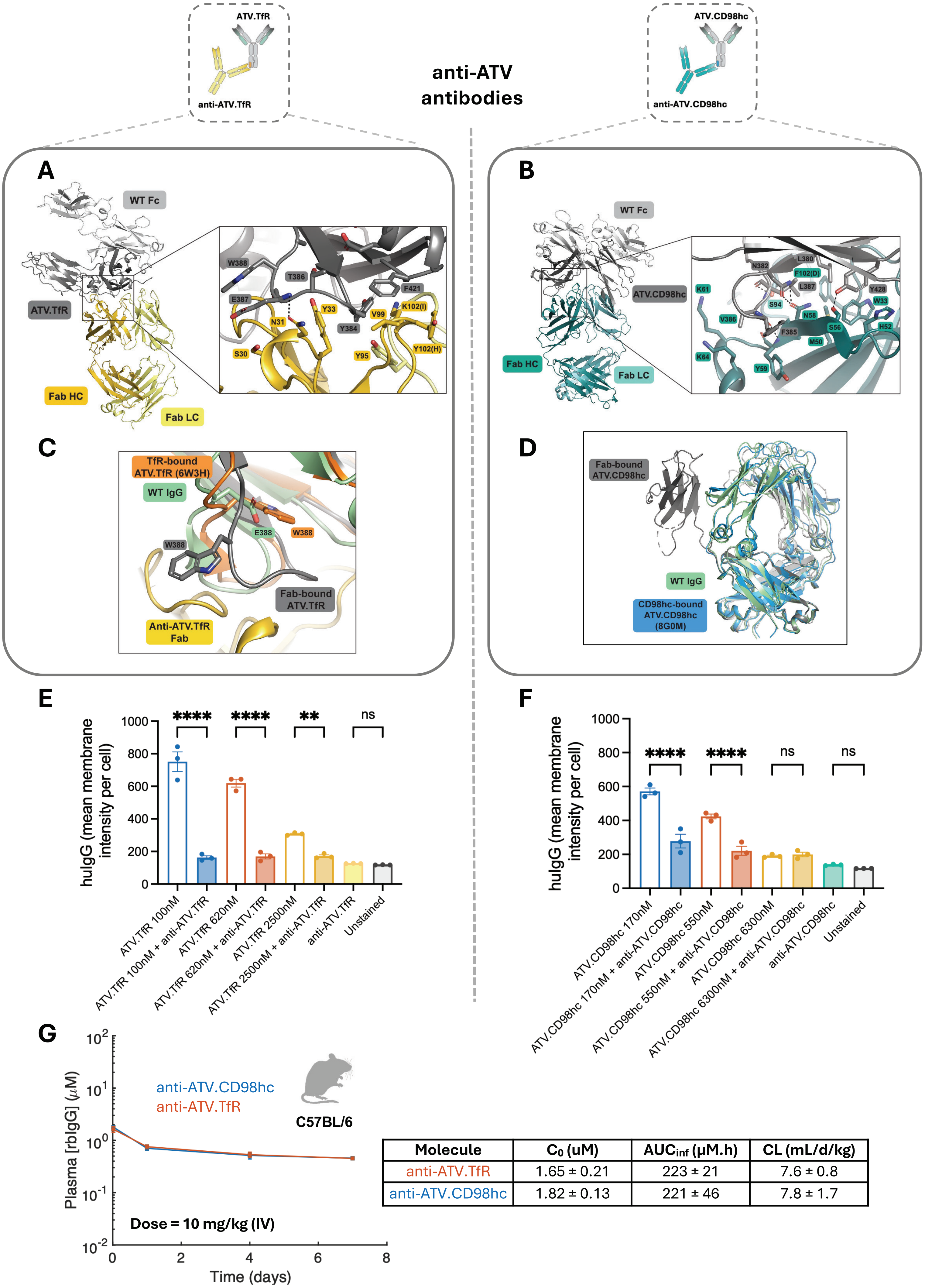


| Molecule | $C_0$ (uM) | $AUC_{inf}$ (μM.h) | CL (mL/d/kg) |
|---|---|---|---|
| anti-ATV.TfR | 1.65 ± 0.21 | 223 ± 21 | 7.6 ± 0.8 |
| anti-ATV.CD98hc | 1.82 ± 0.13 | 221 ± 46 | 7.8 ± 1.7 |

**Fig. S6. Structural, functional, and pharmacokinetic characterization of anti-ATV.TfR and anti-ATV.CD98hc antibodies. (A,B)** Cartoon representations of crystal structures of the complexes of **(A)** anti-ATV.TfR Fab (HC, gold; LC, pale yellow) with Fc-only ATV.TfR (knob, dark gray; hole, light gray) and **(B)** anti-ATV.CD98hc Fab (HC, teal; LC, light teal) with Fc-only ATV.CD98hc (knob, dark gray; hole, light gray)**.** Insets show zoomed in views of Fab-ATV interfaces, with interacting residues shown in sticks. **(C)** Alignment of ATV.TfR from Fab-bound structure (this work, gray) with TfR-bound structure (PDB: 6W3H, orange) and wild type IgG (PDB: 5JII, light green). In wild type IgG and the TfR-bound structure, the residue at position 388 is buried within the CH3 domain. In contrast, in the Fab-bound structure, the CD loop (residues 382-390) of ATV.TfR adopts a unique conformation that allows W388 to interact with the Fab (gold). **(D)** Alignment of ATV.CD98hc from Fab-bound structure (this work, gray) with CD98hc-bound structure (PDB: 8G0M, blue) and wild type IgG (PDB: 5JII, light green). In the Fab-bound structure, the CH2 domain of ATV.CD98hc is displaced from its canonical position, likely due to steric clash from binding of the Fab and/or a potential increase in flexibility between the CH2 and CH3 domains as observed in other engineered Fc constructs (4-7). **(E,F)** HEK293T cells were treated for 1hr at 37 °C with 250 nM **(E)** anti-ATV.TfR or ATV.TfR–anti-ATV.TfR or **(F)** anti-ATV.CD98hc or ATV.CD98hc–anti-ATV.CD98hc. Cell binding and competition were evaluated by immunocytochemistry with high-content imaging. Cells were stained with AlexaFluor®488-conjugated anti-human IgG (1:1000), DAPI, and Deep Red Cell Mask. For each replicate (n=3), ≥20 fields of view were acquired at 40× magnification on the Opera Phenix High-Content Imaging System (PerkinElmer). Images were analyzed using Harmony software (PerkinElmer v4.9). Data are mean ± SD and statistical analysis was performed using one-way ANOVA followed by Tukey's multiple comparison test, *$p < 0.05$, **$p < 0.01$, ***$p < 0.001$, ****$p < 0.0001$, ns: non-significant. **(G).** Rabbit anti-ATV.TfR and anti-ATV.CD98hc antibodies were administered as a single intravenous dose (10 mg/kg) to C57BL/6 mice. Plasma samples were collected up to 168 h post-dose, and antibody concentrations were quantified by MSD. Data represent mean ± SD (n = 5 mice per group). Table represents the key PK parameters of the profiles in panel **(G).**

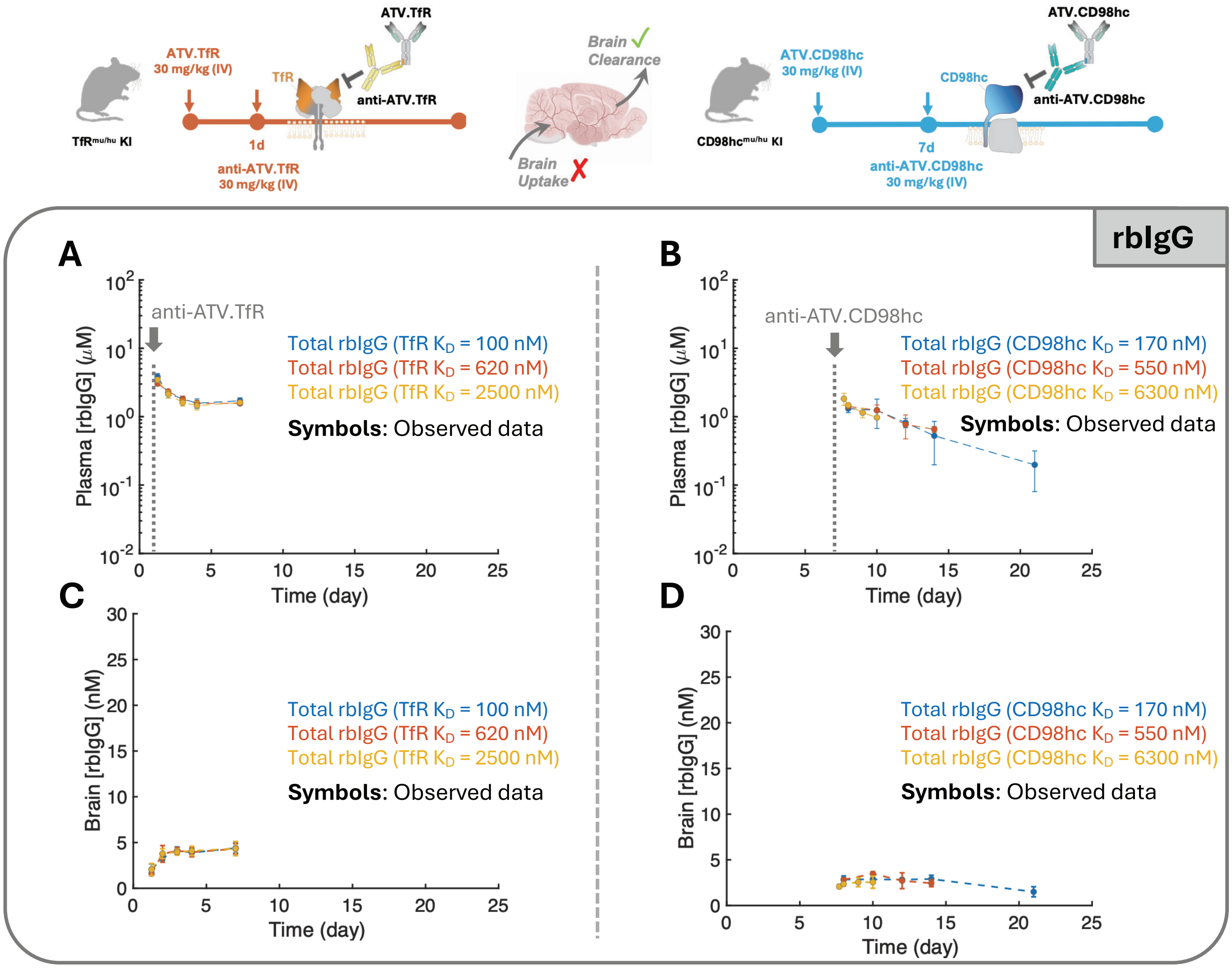

ATV.TfR
ATV.TfR
30 mg/kg (IV)
TfR
anti-ATV.TfR
TfRmu/hu KI
1d
anti-ATV.TfR
30 mg/kg (IV)
Brain Clearance
Brain Uptake
ATV.CD98hc
ATV.CD98hc
30 mg/kg (IV)
CD98hc
anti-ATV.CD98hc
CD98hcmu/hu KI
7d
anti-ATV.CD98hc
30 mg/kg (IV)
rbIgG
A
anti-ATV.TfR
Plasma [rbIgG] (μM)
Time (day)
Total rbIgG (TfR KD = 100 nM)
Total rbIgG (TfR KD = 620 nM)
Total rbIgG (TfR KD = 2500 nM)
Symbols: Observed data
B
anti-ATV.CD98hc
Plasma [rbIgG] (μM)
Time (day)
Total rbIgG (CD98hc KD = 170 nM)
Total rbIgG (CD98hc KD = 550 nM)
Total rbIgG (CD98hc KD = 6300 nM)
Symbols: Observed data
C
Brain [rbIgG] (nM)
Time (day)
Total rbIgG (TfR KD = 100 nM)
Total rbIgG (TfR KD = 620 nM)
Total rbIgG (TfR KD = 2500 nM)
Symbols: Observed data
D
Brain [rbIgG] (nM)
Time (day)
Total rbIgG (CD98hc KD = 170 nM)
Total rbIgG (CD98hc KD = 550 nM)
Total rbIgG (CD98hc KD = 6300 nM)
Symbols: Observed data

**Fig. S7. Observed plasma and brain PK profiles of total rbIgG during pulse–chase experiments.** Observed plasma (A,B) and brain (C,D) concentrations of total rabbit IgG (rbIgG) corresponding to the pulse–chase experiments shown in Fig. 4. $TfR^{mu/hu}$ KI and $CD98hc^{mu/hu}$ KI mice received a single 30 mg/kg intravenous dose of ATV followed by the corresponding anti-ATV administration at day 1 for ATV.TfR and day 7 for ATV.CD98hc. Data are presented as mean ± SD.

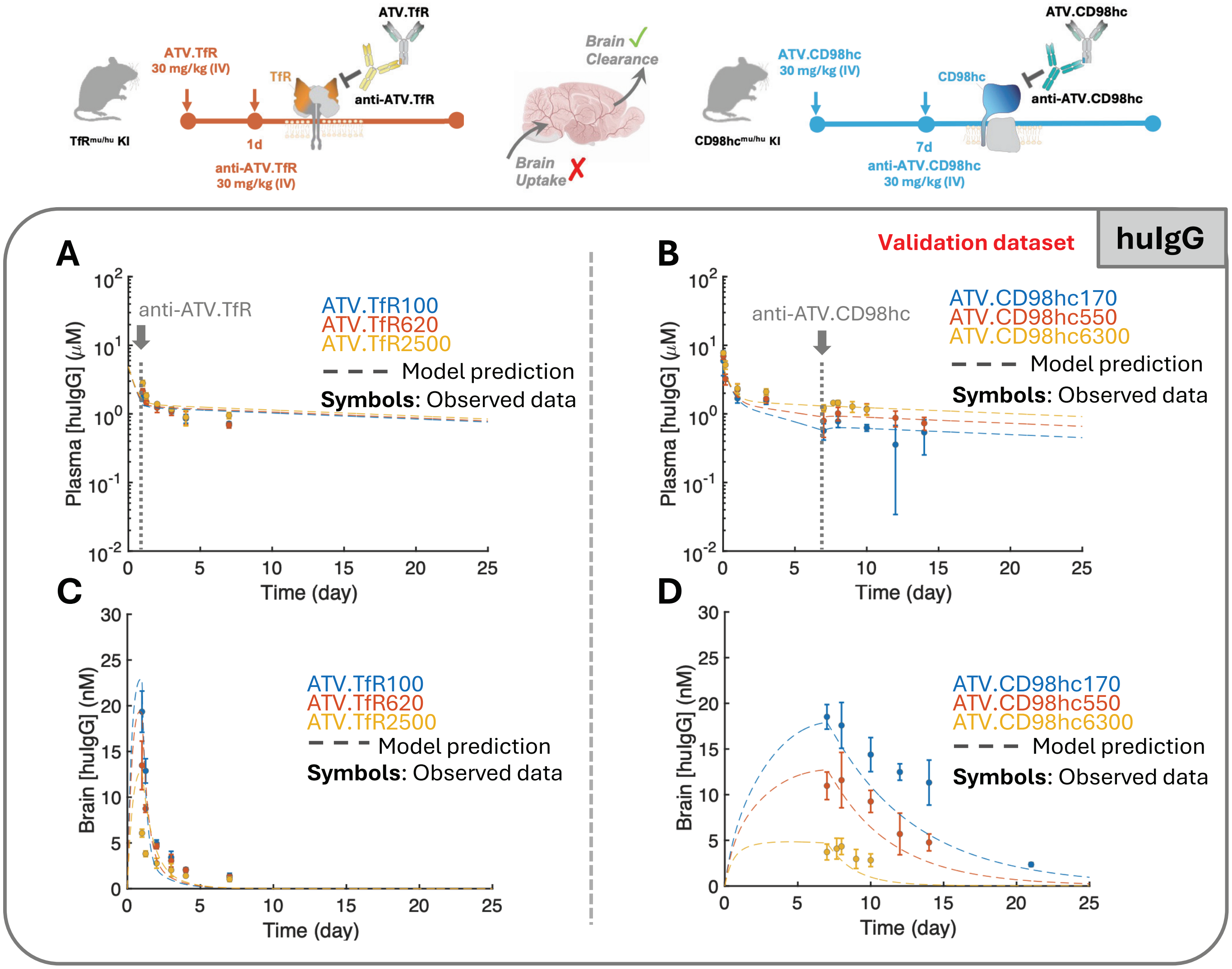
ATV.TfR
ATV.TfR
30 mg/kg (IV)
TfR
anti-ATV.TfR
1d
anti-ATV.TfR
30 mg/kg (IV)
Brain Clearance
Brain Uptake
ATV.CD98hc
ATV.CD98hc
30 mg/kg (IV)
CD98hc
anti-ATV.CD98hc
7d
anti-ATV.CD98hc
30 mg/kg (IV)
Validation dataset
huIgG
A
anti-ATV.TfR
ATV.TfR100
ATV.TfR620
ATV.TfR2500
Model prediction
Symbols: Observed data
Plasma [huIgG] (μM)
Time (day)
B
anti-ATV.CD98hc
ATV.CD98hc170
ATV.CD98hc550
ATV.CD98hc6300
Model prediction
Symbols: Observed data
Plasma [huIgG] (μM)
Time (day)
C
ATV.TfR100
ATV.TfR620
ATV.TfR2500
Model prediction
Symbols: Observed data
Brain [huIgG] (nM)
Time (day)
D
ATV.CD98hc170
ATV.CD98hc550
ATV.CD98hc6300
Model prediction
Symbols: Observed data
Brain [huIgG] (nM)
Time (day)

**Fig. S8. Validation of the calibrated PK model against the pulse–chase experiments.** Experimental data overlaid with model predictions corresponding to the pulse–chase experiments shown in Fig. 4, in which $TfR^{mu/hu}$ KI and $CD98hc^{mu/hu}$ KI mice received a single 30 mg/kg intravenous dose of ATV followed by the corresponding anti-ATV administration at day 1 for ATV.TfR and day 7 for ATV.CD98hc. Plasma (A,B) and brain (C,D) PK profiles of total human IgG (huIgG) corresponding to the ATV.TfR and ATV.CD98hc pulse–chase experiments are shown for each affinity variant. Symbols represent the observed huIgG concentrations, and dashed lines represent predictions from the ATV PK model. Observed data were not used for model calibration and served as an independent validation of the calibrated ATV PK model. Data are presented as mean ± SD.
